# Discovery of the First-in-class G9a/GLP PROTAC Degrader

**DOI:** 10.1101/2024.02.26.582210

**Authors:** Julia Velez, Yulin Han, Hyerin Yim, Peiyi Yang, Zhijie Deng, Kwang-su Park, Md Kabir, H. Ümit Kaniskan, Yan Xiong, Jian Jin

**Author notes:** Corresponding: Jian Jin, Yan Xiong. These authors contributed equally to this work (Julia Velez, Yulin Han).

## Abstract

Aberrantly expressed lysine methyltransferases G9a and GLP, which catalyze mono- and di-methylation of histone H3 lysine 9 (H3K9), have been implicated in numerous cancers. Recent studies have uncovered both catalytic and non-catalytic oncogenic functions of G9a/GLP. As such, G9a/GLP catalytic inhibitors have displayed limited anticancer activity. Here, we report the discovery of the first-in-class G9a/GLP proteolysis targeting chimera (PROTAC) degrader, **10** (MS8709), as a potential anticancer therapeutic. **10** induces G9a/GLP degradation in a concentration-, time, and ubiquitin-proteasome system (UPS)-dependent manner, does not alter the mRNA expression of G9a/GLP and is selective for G9a/GLP over other methyltransferases. Moreover, **10** displays superior cell growth inhibition to the parent G9a/GLP inhibitor UNC0642 in prostate, leukemia, and lung cancer cells and has suitable mouse pharmacokinetic properties for *in vivo* efficacy studies. Overall, **10** is a valuable chemical biology tool to further investigate the functions of G9a/GLP and a potential therapeutic for treating G9a/GLP-dependent cancers.

**Graphical Abstract:** 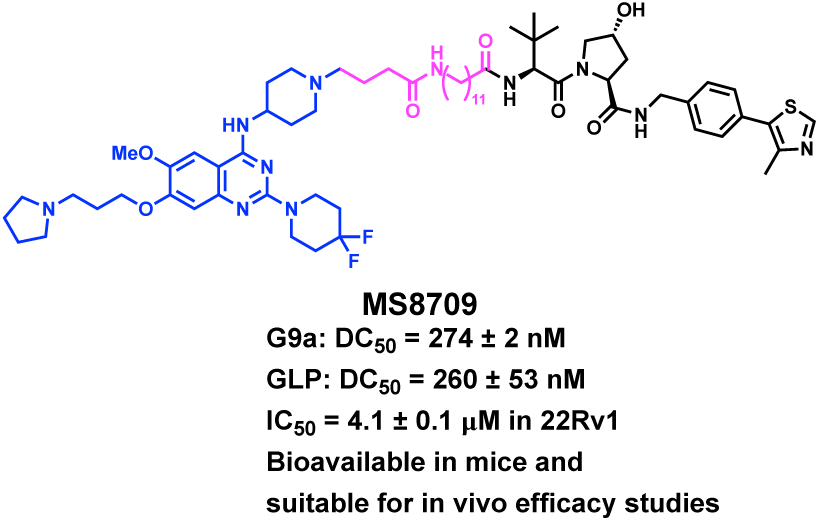

## Introduction

G9a (also known as euchromatic histone-lysine N-methyltransferase 2 (EHMT2)) and G9a-like protein (GLP, also known as EHMT1) are lysine methyltransferases that catalyze mono- and di-methylation of histone H3 lysine 9 (H3K9), transcriptionally repressive chromatin marks, and non-histone proteins.^1–3^ G9a and GLP, which share approximately 80% sequence homology in their catalytic SET domains,^4, 5^ play important roles in diverse cellular processes including cell development, differentiation, and hypoxia response.^6–8^ When atypically expressed, the G9a/GLP axis promotes cancer progression, survival, and metastasis.^2, 9–12^ Overexpression of G9a has been reported in several cancer types such as breast,^13, 14^ lung,^15, 16^ leukemia,^17^ bladder,^18, 19^ colorectal,^20^ and prostate.^21, 22^ G9a/GLP has also been implicated in diseases including sickle cell disease^23^, Prader-Willi syndrome^24^, and Alzheimer’s disease^25^. In addition to G9a/GLP’s well-established histone methyltransferase activity, studies have demonstrated that G9a/GLP have non-canonical oncogenic functions, such as methylating non-histone proteins including HDAC1, DNMT1, and p53.^26–28^ Moreover, it has been shown that G9a/GLP have non-catalytic oncogenic activities by functioning as a co-activator independent of its catalytic domain.^27, 29–33^ For example, G9a positively regulates nuclear receptor gene expression by recruiting GRIP1, CARM1, and p300^27,29, 30^. G9a is also recruited by Runx2 to downstream gene promoters to activate expression^27, 31^, upregulates p21 via interaction with PCAF^32^, and is a co-activator for a subset of p53 downstream genes including pro-apoptotic Puma^33^. As such, enzymatic inhibitors will not target these co-activator functions of G9a/GLP.

Several G9a/GLP enzymatic inhibitors have been reported,^34–44^ including our *in vivo* G9a/GLP chemical probe UNC0642^45^ and G9a/GLP covalent inhibitor MS8511^46^ (**Figure 1**). Though G9a/GLP inhibitors effectively reduce the H3K9me2 mark in cells, they display limited cancer cell killing activity, potentially due to their inability to target G9a/GLP’s non-catalytic oncogenic functions. In contrast, G9a/GLP knockdown led to enhanced anti-proliferative effects through the induction of cell cycle arrest and apoptosis.^8, 47, 48^ Currently, no G9a/GLP inhibitors have progressed to clinic. Therefore, a more effective therapeutic strategy is needed to target G9a/GLP-dependent cancers.

**Figure 1.**
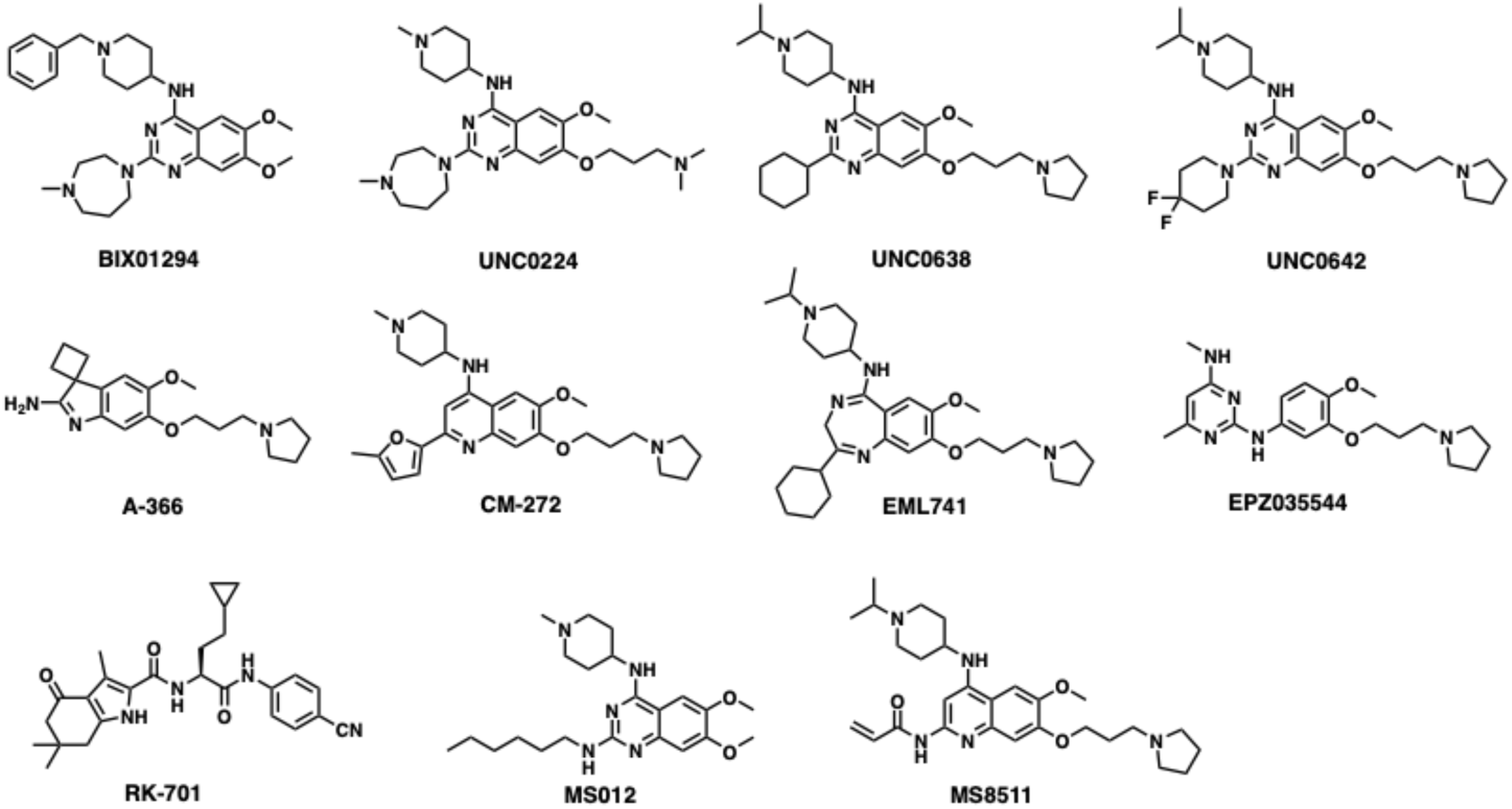
Chemical structures of representative previously reported G9a/GLP catalytic inhibitors.

Proteolysis targeting chimera (PROTAC) degraders are heterobifunctional small molecules that join together a protein of interest (POI) binder and an E3 ubiquitin ligase ligand via a linker. PROTAC induces close proximity between the POI and the E3 ligase, leading to selective polyubiquitination of the POI and its subsequent degradation by the 26S proteasome.^49, 50^ Importantly, PROTAC can eliminate all catalytic and non-catalytic functions by directly degrading the POI. Furthermore, PROTAC has an event-driven pharmacology rather than occupancy-driven pharmacology, making it more efficient to target the POI, and can potentially overcome inhibitor-based resistance.^49, 50^ Thus, PROTAC is an emerging therapeutic modality for the treatment of cancer. Over 20 PROTAC degraders are currently in clinical development.^51, 52^

In this study, we present the discovery of compound **10** (MS8709), the first-in-class G9a/GLP PROTAC degrader, which is based on our previously reported G9a/GLP inhibitor UNC0642 and recruits the von Hippel Lindau (VHL) E3 ligase. Compound **10** potently induces G9a/GLP degradation in prostate, leukemia, and non-small cell lung cancer cells. The G9a/GLP degradation induced by compound **10** is concentration-, time, and ubiquitin-proteasome system (UPS)-dependent, and compound **10** does not alter G9a/GLP transcription. Furthermore, compound **10** is selective for G9a/GLP over other protein methyltransferases. Importantly, compound **10** displays profound anti-proliferative activity and is much more effective than its parent inhibitor, UNC0642. Lastly, compound **10** is bioavailable in mice via intraperitoneal (IP) injection, making it suitable for *in vivo* efficacy studies.

## Results and Discussion

### Design of G9a/GLP PROTAC degraders

Among the previously reported G9a/GLP catalytic inhibitors (Figure 1), we chose UNC0642 as the G9a/GLP binder to generate G9a/GLP PROTAC degraders based on the following reasons. First, UNC0642 is a highly potent G9a/GLP inhibitor (IC50 < 2.5 nM for G9a and GLP in biochemical assays) and more than 300-fold selective for G9a and GLP over a broad range of kinases, GPCRs, transporters, and ion channels.^45^ Second, we previously solved the cocrystal structure of G9a in complex with UNC0638, a close analog of UNC0642 (PDB: 3RJW).^39^ Third, compared to UNC0638, UNC0642 showed improved metabolic stability and in vivo PK properties by replacing the cyclohexyl group with 4,4-difluoropiperidine moiety at 2-position of the quinazoline core.^45^ By analyzing the cocrystal structure mentioned above,^39^ we determined that the *N*-isopropyl moiety was solvent-exposed and could be utilized as a linker attachment point (**Figure 2A**). Our previous structure-activity relationship (SAR) studies revealed that the basicity of this nitrogen was required to maintain high G9a/GLP inhibitory activity.^44^ Based on these results, we replaced the isopropyl moiety with a 4-carbon carboxylic acid to serve as a handle to attach a linker (**Figure 2B**). We then designed a number of putative G9a/GLP PROTAC degraders (**1**-**13**) by conjugating this intermediate to various alkylene and polyethylene glycol (PEG) linkers (**Figure 2B**), which were connected to VHL-1, a classic ligand of the VHL E3 ligase.^53^

**Figure 2.**
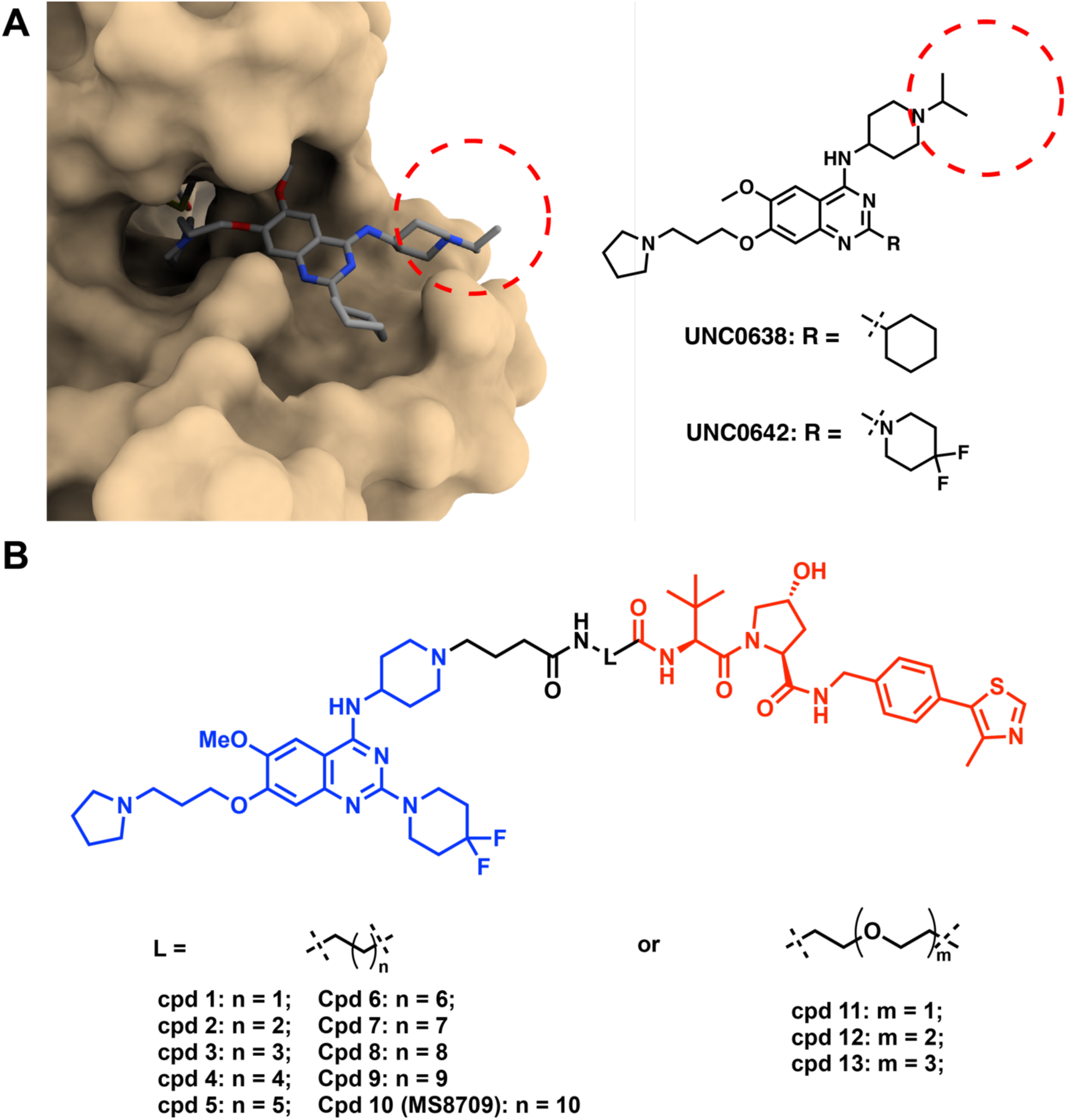
Design of G9a/GLP PROTAC degraders. A) Left, the cocrystal structure of UNC0638 in complex with G9a (PDB ID: 3RJW); Right, chemical structures of UNC0638 and UNC0642. Red dash circle indicates the solvent-exposed region. B) Chemical structures of designed G9a/GLP PROTAC degraders.

### Evaluation of G9a/GLP PROTAC degraders in prostate cancer cells

We first evaluated the putative G9a/GLP PROTAC degraders using western blotting (WB) analysis in the prostate cancer 22Rv1 cell line. The cells were treated with each degrader at 0.3 or 3 μM for 24 h. As shown in **Table 1** and **Figure S1**, compound **1** with a 2-carbon linker induces moderate G9a/GLP degradation, and compounds **2 – 5** (with 3 to 6-carbon linkers, respectively) did not significantly induce G9a/GLP degradation (<30% degradation). Notably, compounds **6 – 10**, which have a longer carbon linker, induced more profound G9a and GLP degradation at 3 μM (>50% degradation). Among them, compounds **8 – 10** induced significant degradation of G9a and GLP at both 0.3 and 3 μM. Most notably, compound **10** (with an 11-carbon linker) induced more than 70% of G9a and around 50% of GLP degradation at 0.3 µM and complete degradation for both proteins at 3 µM (**Table 1** and **Figure S1**). Furthermore, compound **11** (with a 1-PEG linker) did not degrade G9a, but induced significant degradation of GLP at 3 µM (**Table 1** and **Figure S1**). On the other hand, compounds with a longer PEG linker, such as compounds **12** and **13**, induced little or no degradation of G9a/GLP (**Table 1** and **Figure S1**). These results suggest that a relatively long alkyl linker is preferred for G9a/GLP degradation. Because compound **10** is the most effective G9a/GLP degrader among these compounds, we selected compound **10** for further characterization.

**Table 1.** The effect of compounds **1-13** on reducing G9a and GLP protein levels in 22Rv1 cells.^*a*^. ^a^22Rv1 cells were treated with DMSO or the indicated compound at 0.3 or 3 µM for 24 h. The cell lysates were analyzed via WB to examine G9a and GLP protein levels with vinculin as the loading control. The percent of protein degradation was determined by normalizing to DMSO controls. Results shown are the mean values ± SD from two independent experiments. N.D.: no degradation.

| Compound | G9a Degradation (%) |  | GLP Degradation (%) |  |
| --- | --- | --- | --- | --- |
| | 0.3 μM | 3 μM | 0.3 μM | 3 μM |
| 1 | 66 ± 23 | 61 ± 22 | 54 ± 23 | 70 ± 1 |
| 2 | <8 | N.D. | <3 | 21 ± 0.6 |
| 3 | N.D. | <11 | <9 | <11 |
| 4 | N.D. | <30 | <5 | <19 |
| 5 | <10 | <30 | <7 | <20 |
| 6 | N.D. | 75 ± 14 | <28 | 68 ± 9 |
| 7 | N.D. | 87 ± 13 | <27 | 86 ± 2 |
| 8 | 67 ± 22 | <92 | 84 ± 16 | <96 |
| 9 | 79 ± 16 | <92 | 69 ± 15 | <94 |
| 10 | 73 ± 0.1 | >99 | 49 ± 11 | >99 |
| 11 | N.D. | N.D. | <10 | 88 ± 4 |
| 12 | N.D. | <13 | <7 | 21 ± 6 |
| 13 | N.D. | N.D. | <16 | <27 |

### Compound 10 induces G9a/GLP degradation in a concentration- and time-dependent manner and inhibits 22Rv1 cell growth

Next, we characterized the degradation profile of compound **10** in 22Rv1 cells. Compound **10**, but not its parent inhibitor UNC0642, potently induced degradation of G9a and GLP at 1 µM following 24 h treatment (**Figure 3A**). Consistent with its G9a/GLP degradation activity, compound **10** inhibited 22Rv1 cell growth after 7 d treatment with a GI_50_ of 4.1 µM, while UNC0642 did not (**Figure 3B**). This data highlights the superior anti-proliferative activity of G9a/GLP PROTAC degraders to G9a/GLP catalytic inhibitors. Moreover, compound **10** does not affect the cell viability of normal prostate cells (**Figure S2**). Next, we determined that compound **10** induced G9a and GLP degradation in a concentration-dependent manner with the half-maximal degradation concentration (DC_50_) for G9a and GLP as 274 ± 2 and 260 ± 53 nM, respectively (**Figure 3C**). Furthermore, the kinetics of compound **10**-induced G9a/GLP degradation were evaluated in 22Rv1 cells. As shown in Figure 3D, compound **10** induced G9a and GLP degradation in a time-dependent manner. The degradation of G9a and GLP was observed as early as 4 h and reached complete degradation around 24 h, with this effect lasting up to 48 h (**Figure 3D**). Overall, compound **10** potently degrades G9a and GLP in a concentration- and time-dependent manner and has more effective anti-proliferative activity in 22Rv1 cells compared to the parent G9a/GLP catalytic inhibitor UNC0642.

**Figure 3.**
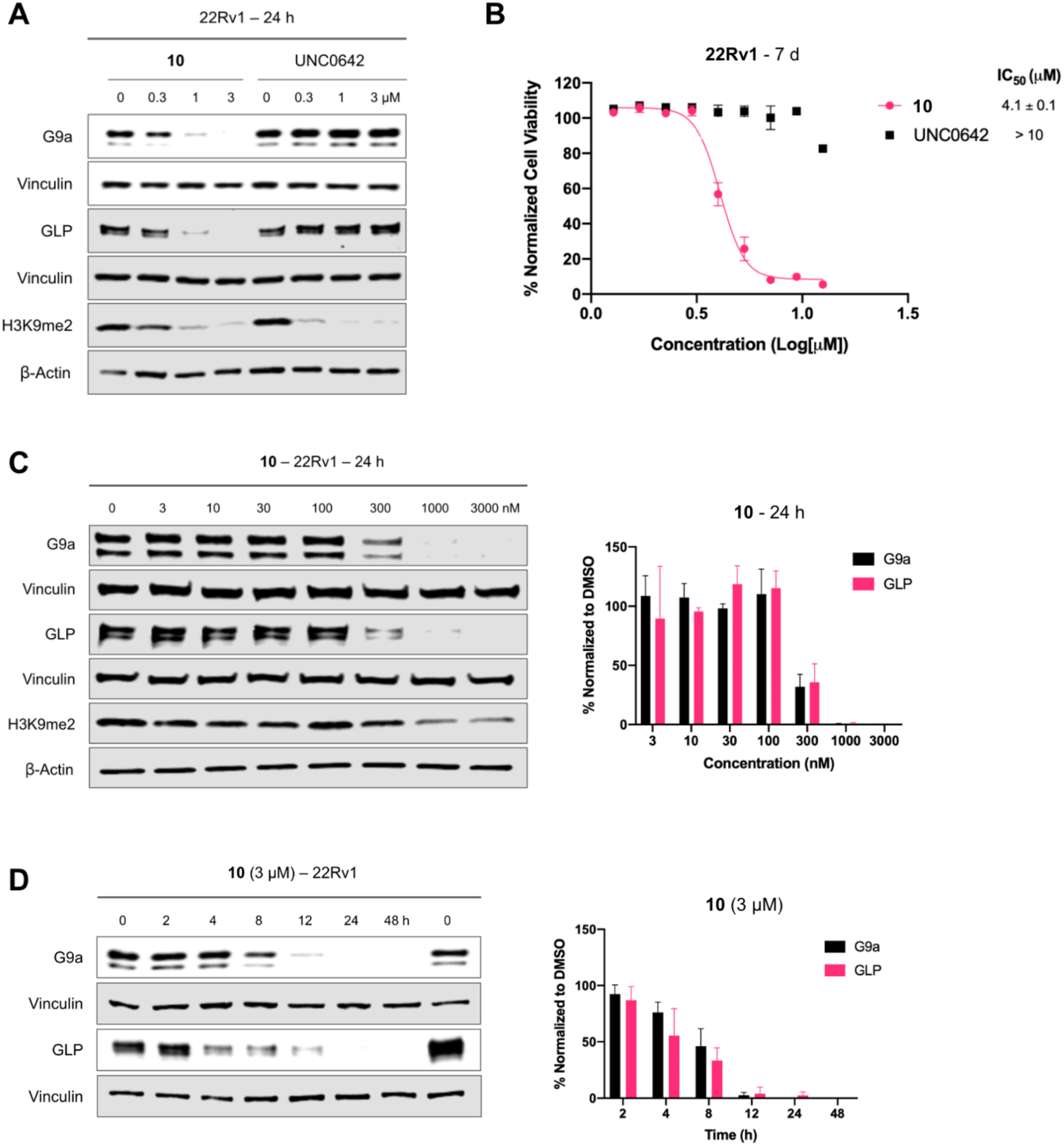
Characterization of compound **10** in 22Rv1 cells. A) WB analysis of G9a, GLP, and H3K9me2 levels following treatment of 22Rv1 cells with **10** or UNC0642 at the indicated concentration (0, 0.3, 1, or 3 μM) for 24 h. The results shown are representative of two independent biological experiments. B) Cell viability of 22Rv1 cells following 7 days treatment with **10** or UNC0642 at the indicated concentration in a WST-8 assay (CCK-8). The mean value ± SD for each concentration point (in technical triplicates from four biological experiments) is shown in the curves. GraphPad Prism 8 was used in analysis of raw data. C) Left, WB analysis of G9a, GLP, and H3K9me2 levels following treatment of 22Rv1 cells with **10** at the indicated concentration for 24 h. Vinculin and β-actin were used at loading controls for WB analysis. The results shown are representative of two independent biological experiments. Right, quantification of relative G9a and GLP protein levels for the WB results shown on the left and its biological repeat. D) Left, WB analysis of G9a and GLP protein levels following treatment of 22Rv1 cells with 3 µM of **10** at the indicated time point. Vinculin was used at the loading control for WB analysis. The results shown are representative of two independent biological experiments. Right, quantification of relative G9a and GLP protein levels for the WB results shown on the left and its biological repeat. Error bars in panels C-D represent ± SD from two independent biological experiments.

#### Compound 10-induced G9a/GLP degradation is VHL- and UPS-dependent

To evaluate the mechanism of action (MOA) of compound **10**-induced G9a/GLP degradation, we first developed compound **14** (MS8709N), a structurally similar analog of compound **10** as a negative control. Compound **14** was designed by incorporating a diastereomer of VHL-1 to block the VHL engagement^54^ while keeping the same G9a/GLP binder and linker (**Figure 4A**). We determined that compound **10**, its parent inhibitor UNC0642, and its negative control **14** all similarly inhibited G9a (**Figure 4B**) and GLP (**Figure 4C**) enzymatic activity, demonstrating that the addition of the linker and VHL ligand did not significantly impact G9a/GLP inhibition. We next assessed the effect of compounds **10** and **14** on G9a and GLP protein levels following 24 h treatment in 22Rv1 cells. As anticipated, compound **10** effectively induced G9a and GLP degradation while **14** did not (**Figure 4D**), indicating that the VHL E3 ligase is crucial for compound **10**-mediated G9a/GLP degradation. Moreover, only compound **10** was able to inhibit 22Rv1 cell colony formation in a clonogenicity assay compared to UNC0642 and compound **14**, suggesting that the G9a/GLP degradation activity is likely the main contributor to the antitumorigenic effect of compound **10** and such antitumorigenic activity is VHL-dependent (**Figure 4E**). To further investigate the MOA of compound **10**, we performed rescue experiments in 22Rv1 cells that were pre-treated with the parent inhibitor (UNC0642, 10 µM), a proteasome inhibitor (MG132, 5 µM), a neddylation inhibitor (MLN4942, 1 µM), or the VHL ligand (VHL-1, 5 µM) for 2 h. The cells were subsequently treated for 24 h with 1 µM of compound **10**. As expected, compound **10**-induced G9a and GLP degradation was rescued upon pre-treatment, indicating that G9a/GLP, VHL, and UPS engagements are necessary for compound **10**-mediated G9a/GLP degradation (**Figure 4F**). Furthermore, compound **10** as well as UNC0642 did not significantly alter the mRNA levels of *G9a* and *GLP* (**Figure S3**), suggesting that compound **10**-induced G9a and GLP degradation is not due to changes in G9a and GLP transcription. Collectively, these results indicate that compound **10** induces G9a/GLP protein degradation in a VHL- and UPS-dependent manner and is a *bona fide* G9a/GLP PROTAC degrader.

**Figure 4.**
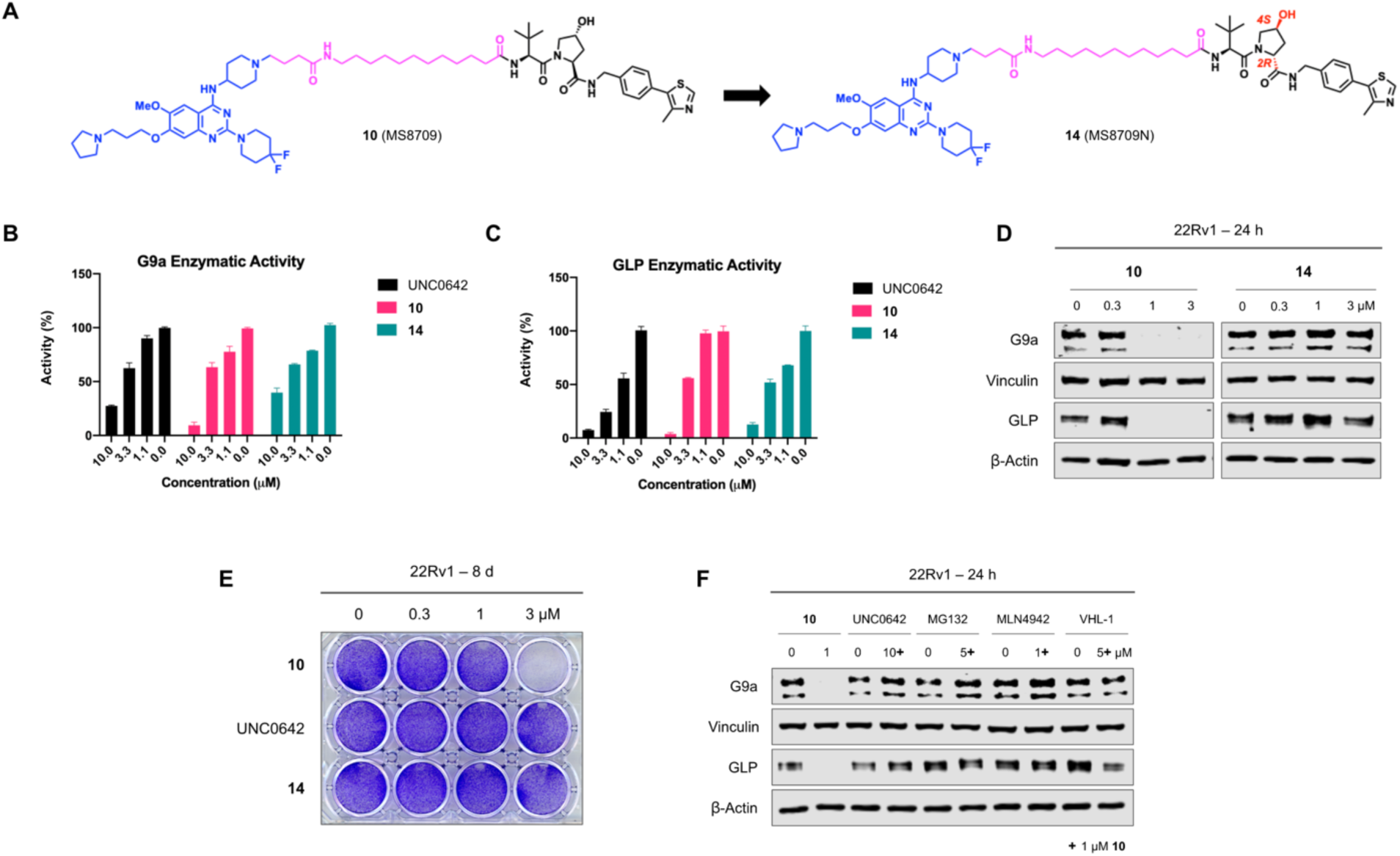
Compound **10** induces G9a/GLP degradation in a VHL- and UPS-dependent manner and compound **10**, but not UNC0642 or its negative control (compound **14**), suppresses clonogenicity in 22Rv1 cells. A) Chemical structures of compound **10** and its negative control, compound **14**, which was designed to abolish binding to VHL. The structural alterations in compound **14** are marked in red. B-C) G9a (B) and GLP (C) enzymatic activity of compound **10**, UNC0642, and compound **14** at indicated concentrations in G9a and GLP radioactive methyltransferase assays. Error bars represent ± SD from two duplicate experiments. D) Representative WB analysis of G9a and GLP protein levels in 22Rv1 cells treated with compound **10** or **14** at the indicated concentration for 24 h. Vinculin and β-actin were used at loading controls. E) Clonogenicity assay of 22Rv1 cells following 8 days treatment with **10**, UNC0642, and **14** at indicated concentrations. Results shown are representative of three independent biological experiments. F) WB analysis of G9a and GLP protein levels in 22Rv1 cells pre-treated with UNC0642 (10 µM), MG132 (5 µM), MLN4942 (1 µM) or VHL-1 (5 µM) for 2 h and subsequently treated with compound **10** (1 µM) for 24 h. Vinculin and β-actin were used as loading controls. WB results shown in panels D and F are representative of two independent biological experiments.

#### Selectivity profile of compound 10

Next, we determined the selectivity of compound **10** against a panel of 21 protein methyltransferases. We did not observe significant inhibition (> 50% at 10 µM) (**Figure 5A**). We also detected the protein levels of EZH2, PRMT7 and SET7/9, which were moderately inhibited by compound **10**, in 22Rv1 cells following the treatment of compound **10**. The WB result indicated compound **10** cannot induce the degradation of these three proteins (**Figure 5B**). Taken together, it suggested that compound **10** is a selective G9a and GLP degrader.

**Figure 5.**
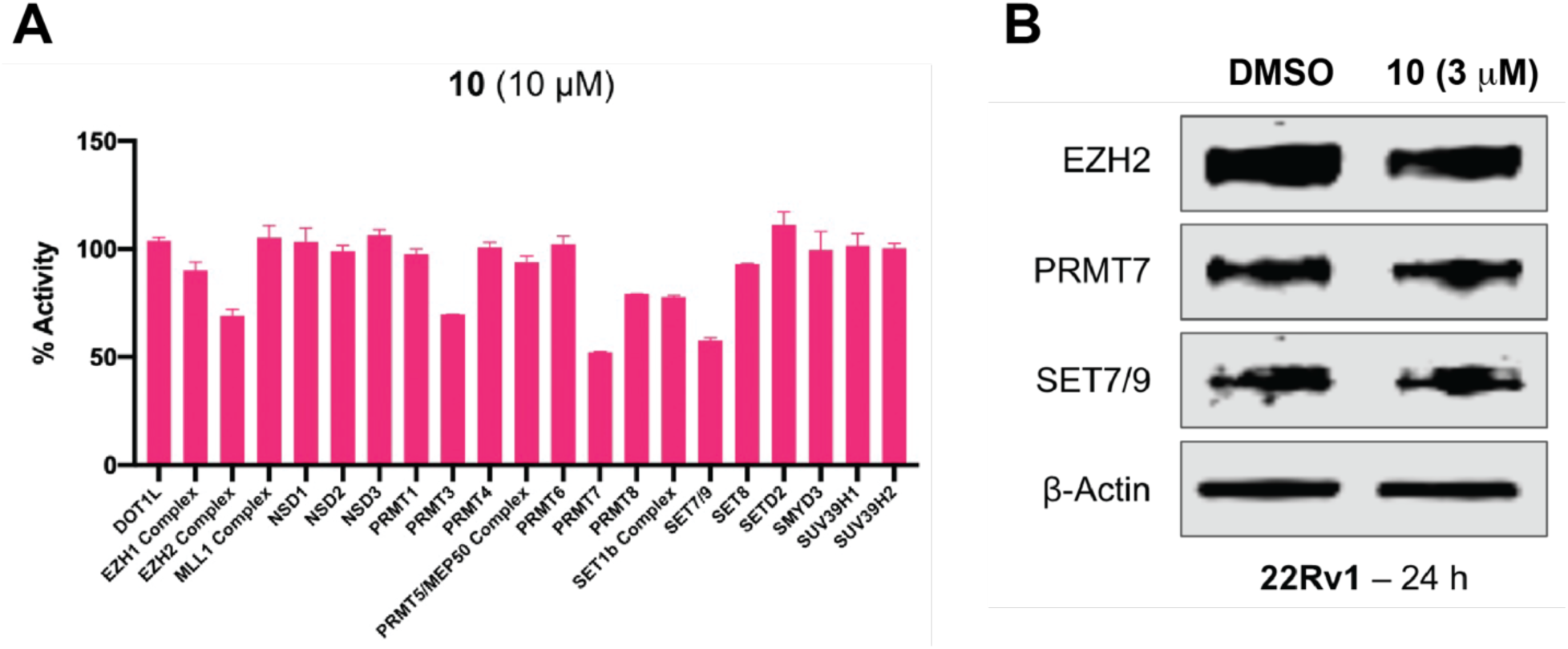
Selectivity profiling of compound **10**. A) Selectivity of compound **10** (at 10 µM) against a panel of 21 protein methyltransferases was assessed using radioactive methyltransferase assays. Results shown are the mean values ± SD from two replicates. B) Representative WB analysis of EZH2, PRMT7 and SET7/9 protein levels in 22Rv1 cells treated with compound **10** at the indicated concentration for 24 h. β-actin was used at the loading control. The WB results shown are representative of two independent experiments.

#### Compound 10 degrades G9a/GLP and inhibits proliferation in leukemia and non-small cell lung cancer cells

We next evaluated effects of compound **10** on G9a/GLP degradation and cell growth inhibition in alternative cancer types, leukemia (using K562 cell line) and non-small cell lung cancer (NSCLC, using H1299 cell line). In K562 cells, compound **10** induced G9a and GLP degradation at 3 µM (**Figure 6A**) and inhibited cell growth with a GI_50_ of 2 ± 0.1 µM following 7 days treatment (**Figure 6B**). On the other hand, UNC0642 did not degrade G9a/GLP (**Figure 6A**) and was unable to suppress cell proliferation (**Figure 6B**). Similarly, compound **10** degraded G9a/GLP at 3 µM in H1299 cells (**Figure 7A**), and inhibited cell growth with a GI_50_ of 5 ± 0.01 µM (**Figure 7B**), while UNC0642 did not. These results indicate that enzymatic inhibition by UNC0642 is insufficient to inhibit the growth of these cancer cells, and the observed cell growth inhibition effect of compound **10** is likely due to its G9a/GLP degradation activity. Overall, compound **10** effectively degrades G9a and GLP and displays superior anti-proliferative activity in K562 and H1299 cells compared to UNC0642.

**Figure 6.**
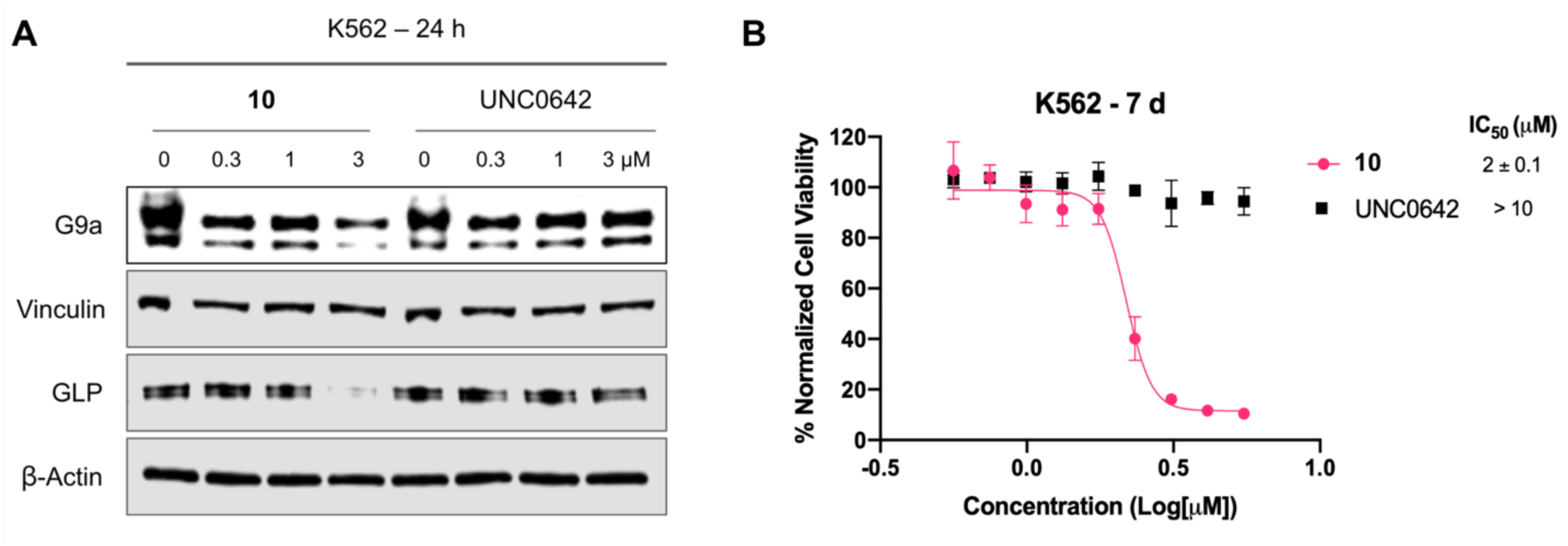
Compound **10** degrades G9a and GLP and inhibits cell growth in K562 cells. A) WB analysis of G9a, GLP, and H3K9me2 levels following treatment of K562 cells with compound **10** or UNC0642 at the indicated concentrations for 24 h. Vinculin and β-actin were used as loading controls. Results shown are representative of two independent biological experiments. B) Cell viability of K562 cells following 7 days treatment with **10** or UNC0642 in a WST-8 assay (CCK-8). Results shown are the mean values ± SD from four independent biological experiments.

**Figure 7.**
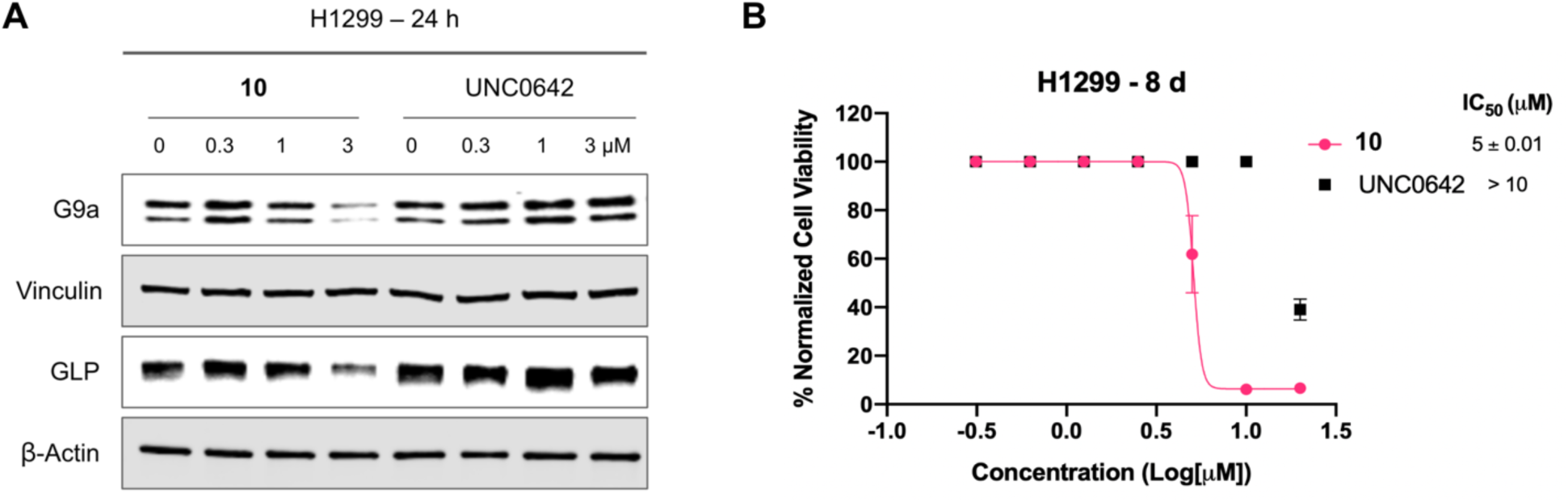
Compound **10** degrades G9a and GLP and inhibits proliferation in H1299 cells. A) WB analysis of G9a and GLP protein levels following treatment of H1299 cells with compound **10** or UNC0642 at the indicated concentration for 24 h. Vinculin and β-actin were used as loading controls. Results shown are representative of two independent biological experiments. B) Cell viability of H1299 cells following 8 days treatment with compound **10** or UNC0642 in a WST-8 assay (CCK-8). Results shown are the mean values ± SD from two independent biological experiments.

#### Compound 10 is bioavailable in mice

Lastly, we evaluated mouse pharmacokinetic (PK) properties of compound **10**. Following a single intraperitoneal (IP) administration of compound **10** at 50 mg/kg dose, compound concentrations in plasma from C57BL6 mice were determined. We were pleased to find that compound **10** showed excellent plasma exposure (**Figure 8** and **Table S1**). It achieved the maximum concentration (C_max_) of 27 ± 0.6 μM at 0.5 h and maintained high plasma exposure levels (> 5 μM) up to 8 h, which are above its G9a/GLP DC_50_ values and GI_50_ values (for cell growth inhibition). In addition, compound **10** was well tolerated by the treated mice, and no clinical signs were observed in the PK study. Taken together, these results suggest that compound **10** has sufficient mouse PK properties and is suitable for *in vivo* efficacy studies.

**Figure 8.**
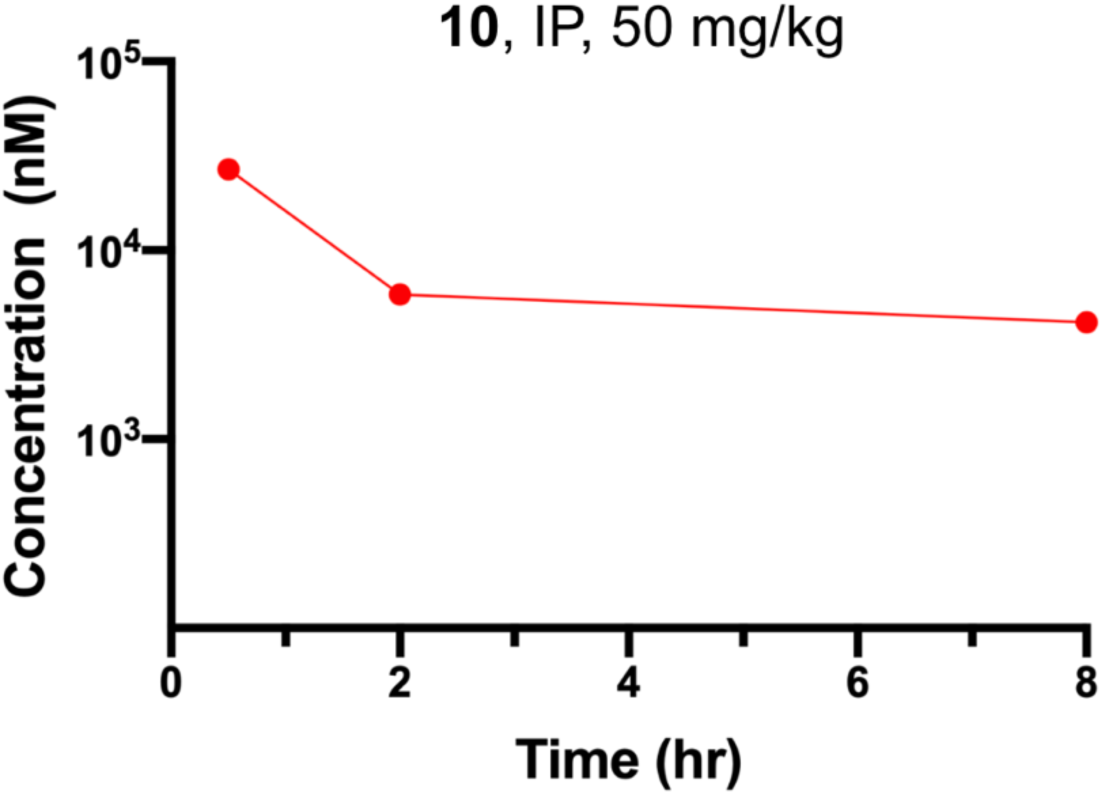
Plasma concentrations of compound **10** over an 8-h period in mice after a single IP injection of compound **10** at 50 mg/kg. The plasma concentrations represent the means ± SEM from three mice per time point.

#### Chemistry synthesis

The synthetic route for compound **1**-**13** is outlined in Scheme **1**. Starting from the previously reported intermediate **I-1**,^45^ the 4-position Cl in the quinazoline ring was first substituted with commercially available intermediate **I-2** under mild condition. The resulting intermediate **I-3** was further installed with the 4,4-difluoropiperidinyl moiety under the microwave irradiation condition. The resulting intermediate **I-4** was then converted into compounds **1**-**13** by coupling with the previously reported VHL linkers **I-5** – **I-17**.^55^

**Scheme 1.**
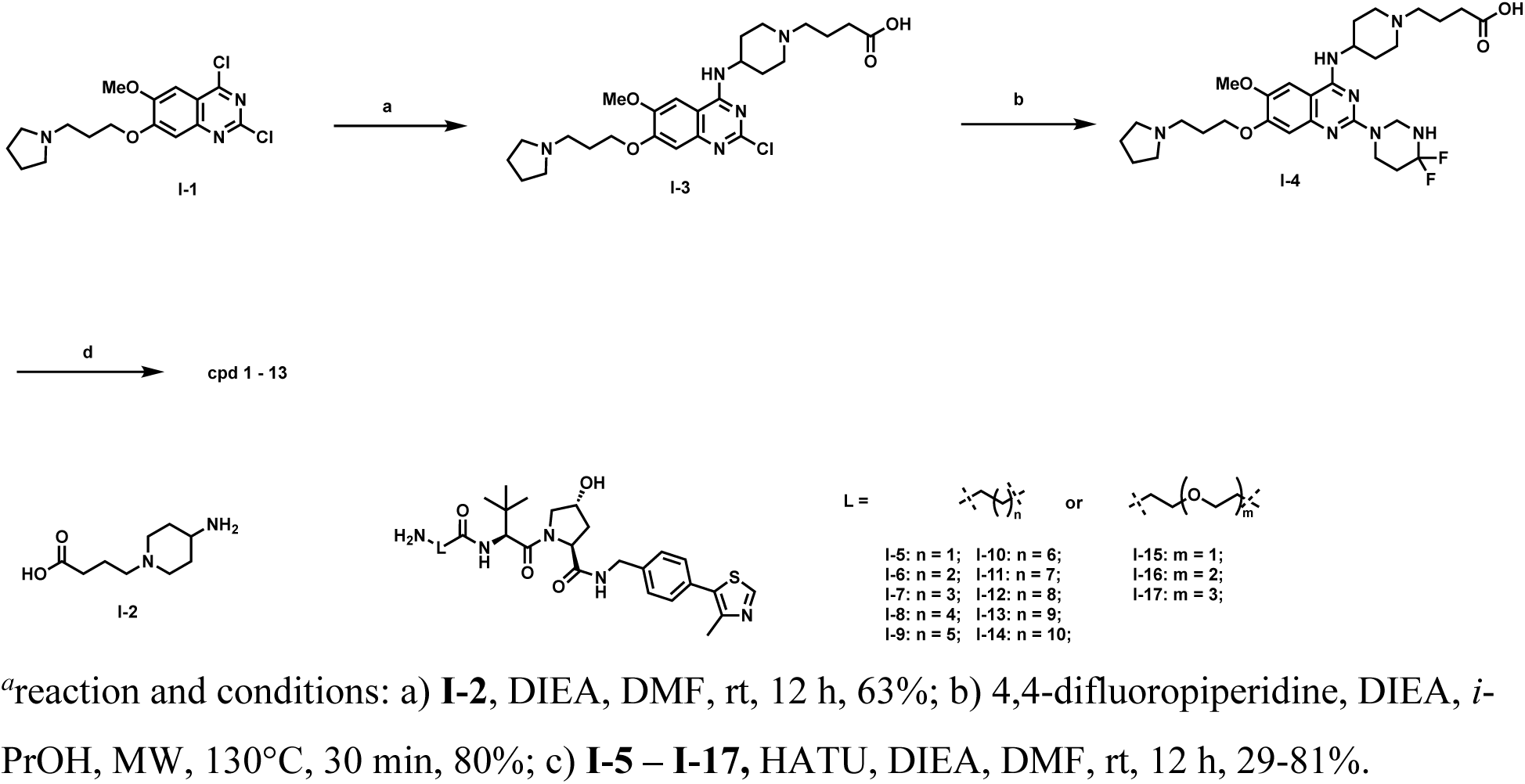
Synthesis of G9a/GLP degraders **1**-**13***^a^* ^a^reaction and conditions: a) **I-2**, DIEA, DMF, rt, 12 h, 63%; b) 4,4-difluoropiperidine, DIEA, *i*-PrOH, MW, 130°C, 30 min, 80%; c) **I-5 – I-17,** HATU, DIEA, DMF, rt, 12 h, 29-81%.

The synthetic route for the negative control compound **14** is straightforward (Scheme **2**). A conventional amide coupling reaction between intermediate **I-4** and intermediate **I-18**, which was prepared from the diastereomer of VHL-1^54^, afforded the negative control compound **14**.

**Scheme 2.**
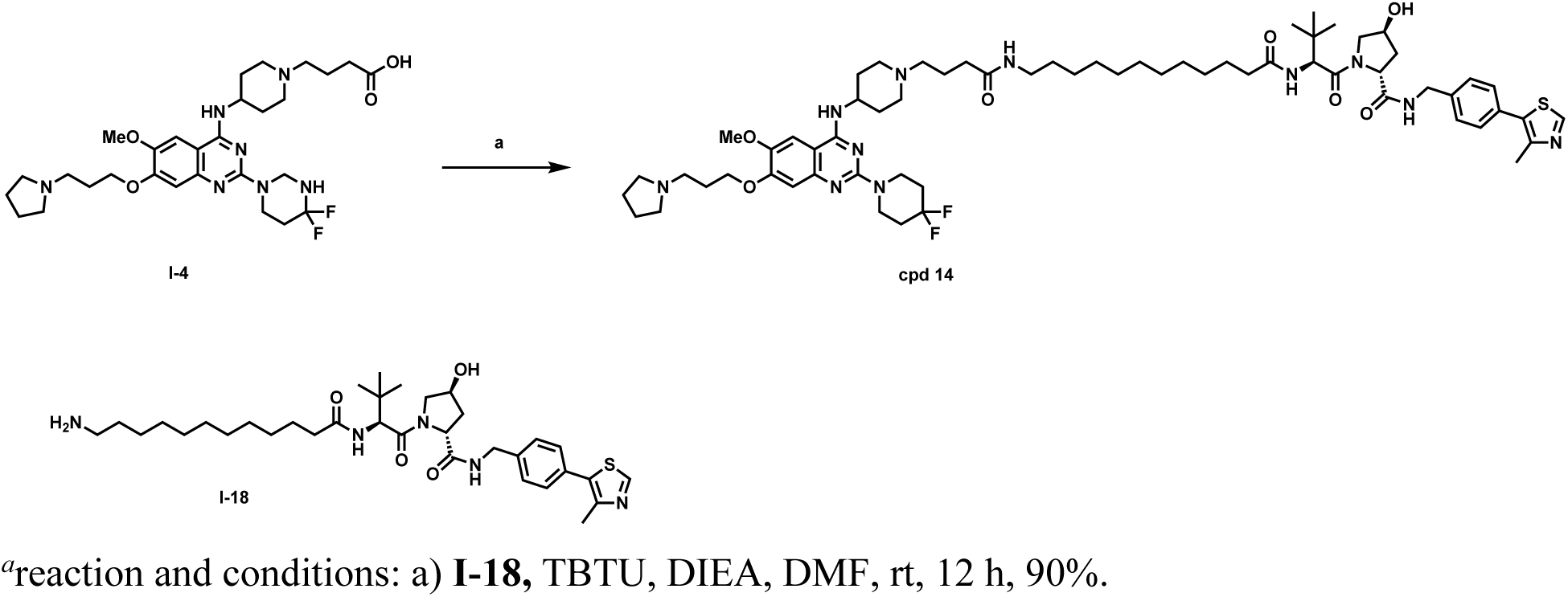
Synthesis of compound **14***^a^* ^a^reaction and conditions: a) **I-18,** TBTU, DIEA, DMF, rt, 12 h, 90%.

## Conclusions

G9a and GLP play crucial roles in various important cellular processes and when abnormally expressed have been implicated in a number of diseases, including cancer. Enzymatic inhibitors of G9a and GLP display limited anti-proliferative activity as they do not target non-catalytic oncogenic functions of G9a and GLP. We employed the PROTAC technology and discovered the first-in-class degrader of G9a and GLP to target both catalytic and non-catalytic oncogenic activities of G9a and GLP. Based on the structural insights revealed from our previously published cocrystal structure of G9a in complex with a catalytic inhibitor, we developed compound **10**, which is based on the G9a/GLP inhibitor UNC0642 and recruits the E3 ligase VHL. Compound **10** induces G9a and GLP degradation in a concentration-, time-, and UPS-dependent manner in prostate cancer 22Rv1 cells. In addition to 22Rv1 cells, compound **10** also degrades G9a and GLP in K562 myelogenous leukemia and H1299 non-small cell lung cancer cell lines. Compound **10**, but not its parent inhibitor UNC0642, suppresses cell growth in 22Rv1, K562, and H1299 cells, demonstrating the superior anti-proliferative activity of G9a/GLP PROTAC degraders to G9a/GLP catalytic inhibitors. Moreover, compound **10** has sufficient mouse PK properties and is suitable for *in vivo* efficacy studies. We also developed compound **14**, a close analog of compound **10**, as a negative control of compound **10**. Overall, we provide compound **10**, which is a novel, potent, and selective PROTAC degrader of G9a and GLP, and its negative control compound **14** to the scientific community as useful chemical biology tools to further study G9a/GLP. Furthermore, compound **10** has potential to be a therapeutic for treating G9a/GLP-dependent cancers.

## Experimental Section

### General chemistry methods

All commercial chemical reagents and solvents were used for the reactions without further purification. Flash column chromatography was performed on Teledyne ISCO CombiFlash Rf+ instrument equipped with a 220/254/280 nm wavelength UV detector and a fraction collector. Normal phase column chromatography was conducted on silica gel columns with either hexane/ethyl acetate or dichloromethane/methanol as eluent. Reverse phase column chromatography was conducted on HP C18 RediSep Rf columns, and the gradient was set to 10% of acetonitrile in H_2_O containing 0.1% TFA progressing to 100% of acetonitrile. All final compounds were purified with preparative high-performance liquid chromatography (HPLC) on an Agilent Prep 1290 infinity II series with the UV detector set to 220/254/280 nm at a flow rate of 40 mL/min. Samples were injected onto a Phenomenex Luna 750 x 30 mm, 5 μm C18 column, and the gradient was set to 10% of acetonitrile in H_2_O containing 0.1% TFA progressing to 100% of acetonitrile. All compounds assessed for biological activity have purity > 95% as determined by an Agilent 1200 series system with DAD detector and a 2.1 mm x 150 mm Zorbax 300SB-C18 5 μm column for chromatography and high-resolution mass spectra (HRMS) that were acquired in positive ion mode using an Agilent G6230BA Accurate Mass TOF with an electrospray ionization (ESI) source. Samples (0.8 μL) were injected onto a C18 column at room temperature, and the flow rate was set to 0.4 mL/min with water containing 0.1% formic acid as solvent A and acetonitrile containing 0.1% formic acid as solvent B. Or acquired in positive ion mode using an Agilent G1969A API-TOF with an electrospray ionization (ESI) source, samples (2 μL) were injected onto a C18 column at room temperature, and the flow rate was set to 0.8 mL/min with water containing 0.1% formic acid as solvent A and acetonitrile containing 0.1% formic acid as solvent B. Nuclear magnetic resonance (NMR) spectra were acquired on Bruker DRX 400 MHz or 600 MHZ for proton (^1^H NMR) and 101 MHz for carbon (^13^C NMR). Chemical shifts for all compounds are reported in parts per million (ppm, δ). The format of chemical shift was reported as follows: chemical shift, multiplicity (s = singlet, d = doublet, t = triplet, q = quartet, m = multiplet), coupling constant (J values in Hz), and integration. All final compounds had > 95% purity using the HPLC methods described above.

**4-(4-((2-chloro-6-methoxy-7-(3-(pyrrolidin-1-yl)propoxy)quinazolin-4-yl)amino)piperidin-1-yl)butanoic acid (I-3).** A suspension of compound **I-1** (prepared according to reported lit^45^) (543.0 mg, 1.5 mmol), 4-(4-aminopiperidin-1-yl)butanoic acid (**I-2**) (commercial available) (777.7 mg, 3.0 mmol), and *N, N*-diisopropylethylamine (1.6 mL, 9.0 mmol) in DMF (30 mL) was stirred overnight at room temperature. Water (10 mL) was added to the reaction mixture and the mixture was extracted with CH_2_Cl_2_. The combined organic layers were concentrated and purified by reversal ISCO to afford the title compound as a yellow solid (816.1 mg, 63%). ^1^H NMR (500 MHz, Methanol-*d*_4_) δ 7.74 (s, 1H), 7.07 (s, 1H), 4.65 – 4.55 (m, 1H), 4.30 (t, *J* = 5.6 Hz, 2H), 3.84 – 3.77 (m, 2H), 3.74 (d, *J* = 11.2 Hz, 2H), 3.46 (t, *J* = 7.2 Hz, 3H), 3.27 – 3.08 (m, 4H), 2.47 (t, *J* = 6.9 Hz, 2H), 2.43 – 2.28 (m, 2H), 2.26 – 2.15 (m, 2H), 2.08 – 2.00 (m, 12H).

**Tert-butyl (3-(4-((2-(4,4-difluoropiperidin-1-yl)-6-methoxy-7-(3-(pyrrolidin-1-yl)propoxy)quinazolin-4-yl)amino)piperidin-1-yl)propyl)carbamate (I-4)** Compound **I-3** (332.7 mg, 0.66 mmol) was dissolved in *i*-PrOH (2 mL) and treated with *N, N*-diisopropylethylamine (0.5 mL, 2.7 mmol) and 4,4-difluoropiperidine salt (212.0 mg, 1.32 mmol). The solution was heated in a microwave reactor at 130 °C for 30 min before being concentrated under reduced pressure. The resulting residue was purified by preparative HPLC to yield title compound as brown solid (310.6 mg, 80%), ^1^H NMR (600 MHz, Methanol-*d*_4_) δ 7.71 (s, 1H), 7.16 (d, *J* = 1.9 Hz, 1H), 4.56 (t, *J* = 11.8 Hz, 1H), 4.26 (t, *J* = 5.5 Hz, 2H), 4.01 (t, *J* = 5.9 Hz, 4H), 3.93 (s, 3H), 3.83 – 3.75 (m, 2H), 3.72 (d, *J* = 12.5 Hz, 2H), 3.44 (t, *J* = 7.2 Hz, 2H), 3.24 – 3.16 (m, 4H), 3.14 – 3.08 (m, 2H), 2.46 (t, *J* = 6.9 Hz, 2H), 2.38-2.26 (m, 4H), 2.22-2.12 (m, 4H), 2.06-1.98 (m, 8H).

Intermediate **I-5** – **I-17** were synthesized following reported lit.^56^

**(2*S*,4*R*)-1-((*S*)-2-(3-(4-(4-((2-(4,4-difluoropiperidin-1-yl)-6-methoxy-7-(3-(pyrrolidin-1-yl)propoxy)quinazolin-4-yl)amino)piperidin-1-yl)butanamido)propanamido)-3,3-dimethylbutanoyl)-4-hydroxy-*N*-(4-(4-methylthiazol-5-yl)benzyl)pyrrolidine-2-carboxamide (cpd 1)** HATU (4.5 mg, 0.011 mmol, 1.0 equiv) and DIPEA (20 µL, 0.22 mmol, 20 equiv) were added to a solution of **I-4** (10.8 mg, 0.011 mmol) and **I-5** (8.2 mg, 1.0 equiv) in DMF (1 mL). After stirring overnight at room temperature, the mixture was subject to preparative HPLC to afford product title product as brown solid (9.7 mg, 62% yield). ^1^H NMR (500 MHz, Methanol-*d*_4_) δ 8.93 (d, *J* = 4.7 Hz, 1H), 7.76 (s, 1H), 7.58 – 7.36 (m, 4H), 7.23 (s, 1H), 4.69 – 4.52 (m, 5H), 4.41 (d, *J* = 15.7, 5.0 Hz, 1H), 4.31 (d, *J* = 5.6 Hz, 2H), 4.12 – 4.03 (m, 4H), 4.02 – 3.94 (m, 3H), 3.89 – 3.73 (m, 5H), 3.56 – 3.44 (m, 4H), 3.29 – 3.12 (m, 6H), 2.63 – 2.34 (m, 12H), 2.33 – 2.00 (m, 14H), 1.08 (s, 9H). HRMS (ESI-TOF) m/z: [M+H]^+^ calcd for C_55_H_78_F_2_N_11_O_7_S^+^ : 1074.5769, Found : 1074.5774.

**Cpds 2** – **13** were synthesized following the same procedure for preparing **cpd 1**.

**((2*S*,4*R*)-1-((*S*)-2-(4-(4-(4-((2-(4,4-difluoropiperidin-1-yl)-6-methoxy-7-(3-(pyrrolidin-1-yl)propoxy)quinazolin-4-yl)amino)piperidin-1-yl)butanamido)butanamido)-3,3-dimethylbutanoyl)-4-hydroxy-*N*-(4-(4-methylthiazol-5-yl)benzyl)pyrrolidine-2-carboxamide)** (**cpd 2**) Brown solid, 62% yield. ^1^H NMR (500 MHz, Methanol-*d*_4_) δ 8.95 (s, 1H), 7.77 (s, 1H), 7.51 – 7.37 (m, 4H), 7.23 (s, 1H), 4.69 – 4.51 (m, 5H), 4.44 – 4.37 (m, 1H), 4.32 (t, *J* = 5.4 Hz, 2H), 4.11 – 4.03 (m, 4H), 3.99 (s, 3H), 3.93 (d, *J* = 11.1 Hz, 1H), 3.90 – 3.74 (m, 5H), 3.50 (t, *J* = 8.5 Hz, 2H), 3.30 – 3.13 (m, 8H), 2.54 – 2.45 (m, 5H), 2.43 – 2.31 (m, 6H), 2.28 – 2.02 (m, 14H), 1.88 – 1.79 (m, 2H), 1.07 (s, 9H). HRMS (ESI-TOF) m/z: [M+H]^+^ calcd for C_56_H_80_F_2_N_11_O_7_S^+^ : 1088.5925, Found : 1088.5933.

**(2S,4R)-1-((S)-2-(5-(4-(4-((2-(4,4-difluoropiperidin-1-yl)-6-methoxy-7-(3-(pyrrolidin-1-yl)propoxy)quinazolin-4-yl)amino)piperidin-1-yl)butanamido)pentanamido)-3,3-dimethylbutanoyl)-4-hydroxy-*N*-(4-(4-methylthiazol-5-yl)benzyl)pyrrolidine-2-carboxamide (cpd 3)** Brown solid, 46% yield. ^1^H NMR (500 MHz, Methanol-*d*_4_) δ 8.93 (s, 1H), 7.77 (s, 1H), 7.62 – 7.41 (m, 4H), 7.22 (s, 1H), 4.71 – 4.46 (m, 5H), 4.40 (d, *J* = 15.5 Hz, 1H), 4.32 (t, *J* = 5.6 Hz, 2H), 4.10 – 4.03 (m, 4H), 3.99 (s, 3H), 3.92 (d, *J* = 10.6 Hz, 1H), 3.88 – 3.81 (m, 3H), 3.78 (d, *J* = 12.4 Hz, 2H), 3.50 (t, *J* = 7.2 Hz, 2H), 3.30 – 3.14 (m, 7H), 2.54 – 2.47 (m, 5H), 2.46 – 2.31 (m, 7H), 2.29 – 2.04 (m, 14H), 1.78 – 1.52 (m, 4H), 1.06 (s, 9H). HRMS (ESI-TOF) m/z: [M+H]^+^ calcd for C_57_H_82_F_2_N_11_O_7_S^+^ : 1102.6082, Found : 1102.6089.

**((2*S*,4*R*)-1-((*S*)-2-(6-(4-(4-((2-(4,4-difluoropiperidin-1-yl)-6-methoxy-7-(3-(pyrrolidin-1-yl)propoxy)quinazolin-4-yl)amino)piperidin-1-yl)butanamido)hexanamido)-3,3-dimethylbutanoyl)-4-hydroxy-*N*-(4-(4-methylthiazol-5-yl)benzyl)pyrrolidine-2-carboxamide) (cpd 4)** Brown solid, 64% yield. ^1^H NMR (500 MHz, Methanol-*d_4_*) δ 8.96 (s, 1H), 7.77 (s, 1H), 7.57 – 7.41 (m, 4H), 7.23 (s, 1H), 4.69 – 4.51 (m, 5H), 4.40 (d, *J* = 15.3 Hz, 1H), 4.32 (t, J = 5.4 Hz, 2H), 4.09 – 4.03 (m, 4H), 3.99 (s, 3H), 3.91 (d, J = 10.6 Hz, 1H), 3.88 – 3.74 (m, 5H), 3.49 (t, J = 7.2 Hz, 2H), 3.29 – 3.14 (m, 6H), 2.55 – 2.44 (m, 5H), 2.45 – 2.02 (m, 22H), 1.72 – 1.52 (m, 2H), 1.61 – 1.53 (m, 2H), 1.47 – 1.36 (m, 2H), 1.06 (s, 9H). HRMS (ESI-TOF) m/z: [M+H]^+^ calcd for C_58_H_84_F_2_N_11_O_7_S^+^ : 1116.6238, Found : 1116.6230.

**(2*S*,4*R*)-1-((*S*)-2-(7-(4-(4-((2-(4,4-difluoropiperidin-1-yl)-6-methoxy-7-(3-(pyrrolidin-1-yl)propoxy)quinazolin-4-yl)amino)piperidin-1-yl)butanamido)heptanamido)-3,3-dimethylbutanoyl)-4-hydroxy-*N*-(4-(4-methylthiazol-5-yl)benzyl)pyrrolidine-2-carboxamide (cpd 5)** Brown solid, 31% yield. ^1^H NMR (500 MHz, Methanol-*d_4_*) δ 8.95 (s, 1H), 7.78 (s, 1H), 7.60 – 7.44 (m, 4H), 7.23 (s, 1H), 4.71 – 4.53 (m, 5H), 4.40 (d, *J* = 15.4 Hz, 1H), 4.33 (t, *J* = 5.6 Hz, 2H), 4.08 (t, *J* = 6.1 Hz, 4H), 4.00 (s, 3H), 3.93 (d, J = 10.9 Hz, 1H), 3.89 –3.74 (m, 5H), 3.51 (t, *J* = 7.2 Hz, 2H), 3.30 – 3.15 (m, 8H), 2.55 – 2.49 (m, 5H), 2.44 – 1.98 (m, 20H), 1.68 – 1.51 (m, 4H), 1.46 – 1.36 (m, 4H), 1.07 (s, 9H). HRMS (ESI-TOF) m/z: [M+H]^+^ calcd for C_59_H_86_F_2_N_11_O_7_S^+^ : 1130.6395, Found : 1130.6414.

**((2*S*,4*R*)-1-((*S*)-2-(8-(4-(4-((2-(4,4-difluoropiperidin-1-yl)-6-methoxy-7-(3-(pyrrolidin-1-yl)propoxy)quinazolin-4-yl)amino)piperidin-1-yl)butanamido)octanamido)-3,3-dimethylbutanoyl)-4-hydroxy-*N*-(4-(4-methylthiazol-5-yl)benzyl)pyrrolidine-2-carboxamide) (cpd 6)** Brown solid, 29% yield. ^1^H NMR (500 MHz, Methanol-*d_4_*) δ 8.94 (s, 1H), 7.78 (s, 1H), 7.59 – 7.41 (m, 4H), 7.23 (s, 1H), 4.72 – 4.50 (m, 5H), 4.40 (d, *J* = 15.1 Hz, 1H), 4.33 (t, *J* = 5.6 Hz, 2H), 4.15 – 4.05 (m, 4H), 4.00 (s, 3H), 3.93 (d, *J* = 10.3 Hz, 1H), 3.89 – 3.75 (m, 5H), 3.50 (t, *J* = 7.2 Hz, 2H), 3.30 – 3.16 (m, 8H), 2.55 – 2.48 (m, 5H), 2.47 – 2.36 (m, 4H), 2.35 – 2.00 (m, 16H), 1.70 – 1.52 (m, 4H), 1.45 – 1.33 (m, 6H), 1.07 (s, 9H). HRMS (ESI-TOF) m/z: [M+H]^+^ calcd for C_60_H_88_F_2_N_11_O_7_S^+^ : 1144.6551, Found : 1144.6570.

**(2*S*,4*R*)-1-((*S*)-2-(9-(4-(4-((2-(4,4-difluoropiperidin-1-yl)-6-methoxy-7-(3-(pyrrolidin-1-yl)propoxy)quinazolin-4-yl)amino)piperidin-1-yl)butanamido)nonanamido)-3,3-dimethylbutanoyl)-4-hydroxy-*N*-(4-(4-methylthiazol-5-yl)benzyl)pyrrolidine-2-carboxamide (cpd 7)** Brown solid, 61% yield. ^1^H NMR (500 MHz, Methanol-*d*_4_) δ 8.95 (s, 1H), 7.78 (s, 1H), 7.57 – 7.42 (m, 4H), 7.23 (s, 1H), 4.74 – 4.51 (m, 5H), 4.40 (d, *J* = 15.5 Hz, 1H), 4.33 (t, *J* = 5.6 Hz, 2H), 4.12 – 4.07 (m, 4H), 4.00 (s, 3H), 3.93 (d, *J* = 10.9 Hz, 1H), 3.90 – 3.75 (m, 5H), 3.51 (t, *J* = 7.2 Hz, 2H), 3.31 – 3.17 (m, 8H), 2.55 – 2.47 (m, 5H), 2.45 – 2.04 (m, 20H), 1.72 – 1.48 (m, 4H), 1.43 – 1.33 (m, 8H), 1.07 (s, 9H). HRMS (ESI-TOF) m/z: [M+H]^+^ calcd for C_61_H_90_F_2_N_11_O_7_S^+^ : 1158.6708, Found : 1158.6717.

**((2*S*,4*R*)-1-((*S*)-2-(10-(4-(4-((2-(4,4-difluoropiperidin-1-yl)-6-methoxy-7-(3-(pyrrolidin-1-yl)propoxy)quinazolin-4-yl)amino)piperidin-1-yl)butanamido)decanamido)-3,3-dimethylbutanoyl)-4-hydroxy-*N*-(4-(4-methylthiazol-5-yl)benzyl)pyrrolidine-2-carboxamide) (cpd 8)** Brown solid, 74% yield. ^1^H NMR (500 MHz, Methanol-*d_4_*) δ 8.95 (s, 1H), 7.76 (s, 1H), 7.53 – 7.40 (m, 4H), 7.22 (s, 1H), 4.70 – 4.51 (m, 5H), 4.39 (d, *J* = 15.5 Hz, 1H), 4.31 (t, *J* = 5.6 Hz, 2H), 4.07 (t, *J* = 5.9 Hz, 4H), 3.98 (s, 3H), 3.92 (d, *J* = 11.0 Hz, 1H), 3.87 – 3.73 (m, 5H), 3.49 (t, *J* = 7.2 Hz, 2H), 3.29 – 3.12 (m, 8H), 2.53 – 2.45 (m, 5H), 2.44 – 2.00 (m, 20H), 1.69 – 1.48 (m, 4H), 1.40 – 1.30 (m, 10H), 1.05 (s, 9H). HRMS (ESI-TOF) m/z: [M+H]^+^ calcd for C_61_H_90_F_2_N_11_O_7_S^+^ : 1158.6708, Found : 1158.6717. HRMS (ESI-TOF) m/z: [M+H]^+^ calcd for C_62_H_92_F_2_N_11_O_7_S^+^ : 1172.6864, Found : 1172.6869.

**((2*S*,4*R*)-1-((*S*)-2-(11-(4-(4-((2-(4,4-difluoropiperidin-1-yl)-6-methoxy-7-(3-(pyrrolidin-1-yl)propoxy)quinazolin-4-yl)amino)piperidin-1-yl)butanamido)undecanamido)-3,3-dimethylbutanoyl)-4-hydroxy-*N*-(4-(4-methylthiazol-5-yl)benzyl)pyrrolidine-2-carboxamide) (cpd 9)** Brown solid, 81% yield. ^1^H NMR (500 MHz, Methanol-*d_4_*) δ 8.96 (s, 1H), 7.76 (s, 1H), 7.57 – 7.34 (m, 4H), 7.22 (s, 1H), 4.71 – 4.50 (m, 5H), 4.38 (d, *J* = 15.5 Hz, 1H), 4.31 (t, *J* = 5.6 Hz, 2H), 4.06 (t, *J* = 5.6 Hz, 4H), 3.98 (s, 3H), 3.92 (d, *J* = 10.9 Hz, 1H), 3.88 – 3.72 (m, 5H), 3.49 (t, *J* = 9.0 Hz, 2H), 3.27 – 3.13 (m, 8H), 2.55 – 2.45 (m, 5H), 2.44 – 2.02 (m, 20H), 1.68 – 1.47 (m, 4H), 1.42 – 1.28 (m, 12H), 1.05 (s, 9H). HRMS (ESI-TOF) m/z: [M+H]^+^ calcd for C_63_H_94_F_2_N_11_O_7_S^+^ : 1186.7021, Found : 1186.7039.

**(2S,4R)-1-((S)-2-(12-(4-(4-((2-(4,4-difluoropiperidin-1-yl)-6-methoxy-7-(3-(pyrrolidin-1-yl)propoxy)quinazolin-4-yl)amino)piperidin-1-yl)butanamido)dodecanamido)-3,3-dimethylbutanoyl)-4-hydroxy-N-(4-(4-methylthiazol-5-yl)benzyl)pyrrolidine-2-carboxamide (cpd 10)** Brown solid, 61% yield. ^1^H NMR (400 MHz, Methanol-*d*_4_) δ 8.94 (s, 1H), 7.74 (s, 1H), 7.47 (d, *J* = 8.0 Hz, 2H), 7.41 (d, *J* = 8.1 Hz, 2H), 7.19 (s, 1H), 4.67 – 4.22 (m, 9H), 4.08 – 3.87 (m, 8H), 3.81 – 3.72 (m, 5H), 3.47 (t, *J* = 7.0 Hz, 2H), 3.27 – 3.11 (m, 8H), 2.48 – 2.43 (m, 5H), 2.39 – 2.35 (m, 4H), 2.28 – 2.12 (m, 9H), 2.12 – 1.99 (m, 7H), 1.65 – 1.45 (m, 4H), 1.39 – 1.23 (m, 13H), 1.04 (s, 9H). ^13^C NMR (101 MHz, Methanol-*d*_4_) δ 176.08, 174.75, 174.51, 172.33, 160.45, 156.15, 153.09, 152.66, 149.28, 148.68, 140.40, 137.22, 131.30, 130.34, 128.97, 104.94, 104.14, 100.74, 71.09, 68.07, 60.86, 59.00, 58.25, 58.03, 57.02, 55.41, 54.25, 52.84, 43.66, 43.38, 43.33, 43.28, 40.68, 38.98, 36.65, 36.59, 34.87, 34.63, 34.40, 34.11, 30.71, 30.64, 30.49, 30.47, 30.36, 30.32, 29.89, 28.07, 27.22, 27.06, 26.60, 23.99, 21.18, 15.75. HRMS (ESI-TOF) m/z: [M+H]^+^ calcd for C_64_H_96_F_2_N_11_O_7_S^+^ : 1200.7177, Found : 1200.7195.

**(2*S*,4*R*)-1-((*S*)-2-(3-(2-(4-(4-((2-(4,4-difluoropiperidin-1-yl)-6-methoxy-7-(3-(pyrrolidin-1-yl)propoxy)quinazolin-4-yl)amino)piperidin-1-yl)butanamido)ethoxy)propanamido)-3,3-dimethylbutanoyl)-4-hydroxy-*N*-(4-(4-methylthiazol-5-yl)benzyl)pyrrolidine-2-carboxamide (cpd 11)** Brown solid, 60% yield. ^1^H NMR (500 MHz, Methanol-*d*_4_) δ 8.94 (s, 1H), 7.77 (s, 1H), 7.60 – 7.39 (m, 4H), 7.21 (s, 1H), 4.71 (t, *J* = 4.6 Hz, 2H), 4.66 – 4.59 (m, 3H), 4.58 – 4.53 (m, 2H), 4.43 (d, *J* = 15.6 Hz, 1H), 4.33 (t, *J* = 5.5 Hz, 2H), 4.11 – 4.05 (m, 4H), 4.00 (s, 3H), 3.94 (d, *J* = 11.0 Hz, 1H), 3.90 – 3.82 (m, 3H), 3.82 – 3.73 (m, 4H), 3.60 (t, *J* = 5.3 Hz, 2H), 3.51 (t, *J* = 7.2 Hz, 2H), 3.45 (t, *J* = 5.3 Hz, 2H), 3.31 – 3.14 (m, 6H), 2.62 – 2.55 (m, 2H), 2.52 (s, 3H), 2.46 – 2.35 (m, 4H), 2.33 – 2.17 (m, 6H), 2.16 – 2.00 (m, 8H), 1.09 (d, *J* = 2.1 Hz, 9H). HRMS (ESI-TOF) m/z: [M+H]^+^ calcd for C_57_H_82_F_2_N_11_O_8_S^+^ : 1118.6031, Found : 1118.6027.

**(2*S*,4*R*)-1-((*S*)-2-(*tert*-butyl)-17-(4-((2-(4,4-difluoropiperidin-1-yl)-6-methoxy-7-(3-(pyrrolidin-1-yl)propoxy)quinazolin-4-yl)amino)piperidin-1-yl)-4,14-dioxo-7,10-dioxa-3,13-diazaheptadecanoyl)-4-hydroxy-*N*-(4-(4-methylthiazol-5-yl)benzyl)pyrrolidine-2-carboxamide (cpd 12)** Brown solid, 35% yield. ^1^H NMR (500 MHz, Methanol-*d*_4_) δ 8.98 (s, 1H), 7.74 (s, 1H), 7.67 (d, *J* = 9.2 Hz, 1H), 7.55 – 7.37 (m, 4H), 7.21 (s, 1H), 4.74 – 4.69 (m, 2H), 4.62 – 4.55 (m, 2H), 4.54 – 4.48 (m, 2H), 4.48 – 4.35 (m, 1H), 4.31 (t, *J* = 5.6 Hz, 2H), 4.14 – 4.03 (m, 4H), 3.97 (s, 3H), 3.94 – 3.87 (m, 1H), 3.88 – 3.80 (m, 3H), 3.78 – 3.60 (m, 10H), 3.56 (t, *J* = 5.5 Hz, 2H), 3.49 (t, *J* = 7.2 Hz, 2H), 3.40 (t, *J* = 5.6 Hz, 2H), 3.27 – 3.11 (m, 5H), 3.00 (s, 1H), 2.55 – 2.46 (m, 5H), 2.44 – 2.32 (m, 4H), 2.29 – 2.01 (m, 14H), 1.06 (s, 9H). HRMS (ESI-TOF) m/z: [M+H]^+^ calcd for C_59_H_86_F_2_N_11_O_9_S^+^ : 1162.6293, Found : 1162.6301.

**(2*S*,4*R*)-1-((*S*)-2-(*tert*-butyl)-20-(4-((2-(4,4-difluoropiperidin-1-yl)-6-methoxy-7-(3-(pyrrolidin-1-yl)propoxy)quinazolin-4-yl)amino)piperidin-1-yl)-4,17-dioxo-7,10,13-trioxa-3,16-diazaicosanoyl)-4-hydroxy-*N*-(4-(4-methylthiazol-5-yl)benzyl)pyrrolidine-2-carboxamide (cpd 13)** Brown solid, 66% yield. ^1^H NMR (500 MHz, Methanol-*d*_4_) δ 9.00 (s, 1H), 7.74 (s, 1H), 7.53 – 7.35 (m, 4H), 7.22 (s, 1H), 4.71 – 4.49 (m, 5H), 4.43 – 4.36 (m, 1H), 4.32 (t,*J* = 5.5 Hz, 2H), 4.07 (t, *J* = 6.0 Hz, 4H), 4.01 – 3.95 (m, 3H), 3.91 (d, *J* = 11.1 Hz, 1H), 3.87 – 3.80 (m, 3H), 3.80 – 3.74 (m, 4H), 3.70 – 3.61 (m, 9H), 3.58 (t, *J* = 5.4 Hz, 2H), 3.50 (t, *J* = 7.0 Hz, 2H), 3.42 (t, *J* = 5.3 Hz, 2H), 3.29 – 3.14 (m, 5H), 3.08 – 2.96 (m, 1H), 2.66 – 2.57 (m, 1H), 2.55 – 2.46 (m, 5H), 2.45 – 2.33 (m, 4H), 2.29 – 2.02 (m, 14H), 1.07 (s, 9H). HRMS (ESI-TOF) m/z: [M+H]^+^ calcd for C_61_H_90_F_2_N_11_O_10_S^+^ : 1206.6555, Found : 1206.6539.

**(2*R*,4*S*)-1-((S)-2-(12-(4-(4-((2-(4,4-difluoropiperidin-1-yl)-6-methoxy-7-(3-(pyrrolidin-1-yl)propoxy)quinazolin-4-yl)amino)piperidin-1-yl)butanamido)dodecanamido)-3,3-dimethylbutanoyl)-4-hydroxy-N-(4-(4-methylthiazol-5-yl)benzyl)pyrrolidine-2-carboxamide (Cpd 14)** TBTU (3.9 mg, 0.012 mmol, 1.2 equiv) and DIPEA (10 µL, 0.1 mmol, 10 equiv) were added to a solution of **I-4** (9.5 mg, 0.01 mmol) and **I-18** (7.7 mg, 1.0 equiv) in DMF (1 mL). After stirring overnight at room temperature, the mixture was subject to preparative HPLC to afford title product as brown solid (14.2 mg, 90% yield). ^1^H NMR (400 MHz, Methanol-*d*_4_) δ 8.91 (s, 1H), 7.73 (s, 1H), 7.43 (d, *J* = 8.1 Hz, 2H), 7.38 (d, *J* = 8.1 Hz, 2H), 7.18 (s, 1H), 4.67 – 4.19 (m, 9H), 4.09 – 3.89 (m, 8H), 3.88 – 3.68 (m, 5H), 3.46 (t, *J* = 7.1 Hz, 2H), 3.25 – 3.11 (m, 8H), 2.49 – 2.45 (m, 5H), 2.40 – 2.33 (m, 4H), 2.27 – 2.13 (m, 9H), 2.11 – 1.96 (m, 7H), 1.49 (t, *J* = 4.1 Hz, 2H), 1.30 – 1.24 (m, 15H), 1.08 (s, 9H). ^13^C NMR (101 MHz, Methanol-*d*_4_) δ 177.04, 174.85, 174.55, 172.61, 160.47, 156.20, 152.95, 152.69, 149.31, 148.89, 140.28, 137.23, 131.37, 130.39, 128.66, 104.93, 104.15, 100.72, 70.53, 68.08, 60.99, 60.15, 58.36, 57.02, 56.83, 55.44, 54.27, 52.89, 43.48, 43.38, 43.34, 43.28, 40.72, 39.13, 36.26, 35.34, 34.89, 34.65, 34.41, 34.18, 30.72, 30.63, 30.48, 30.47, 30.42, 30.34, 29.93, 28.09, 27.06, 26.81, 26.63, 24.01, 21.16, 16.00. HRMS (ESI-TOF) m/z: [M+H]^+^ calcd for C_64_H_96_F_2_N_11_O_7_S^+^ : 1200.7177, Found : 1200.7148.

### Cell culture

22Rv1, H1299, and K562 cells were purchased from ATCC. 22Rv1 and H1299 were cultured in RPMI-1640 medium containing 10% FBS and 1% penicillin-streptomycin. K562 cells were grown in IMDM containing 10% FBS and 1% penicillin-streptomycin. All cells were cultured at 37° C with 5% CO_2_.

### Western blot assay

Cells were lysed with RIPA buffer containing 1x protease-proteasome inhibitor cocktail for 30 minutes. Samples were then centrifuged at 4° C for 12 minutes at 15000 RPM. 1x Laemmli buffer was added to the cell lysates and then heated for 10 minutes at 100° C. Protein concentration was quantified using the Pierce Gold Standard BCA kit (Thermofisher). 10-30 μg of samples were run on 4-15% (BioRad), 4-20% (BioRad), or 6% (Thermofisher) tris-glycine SDS-PAGE gels then transferred onto PVDF membranes using the Transblot rapid transfer machine (BioRad). Following transfer, the membranes were blocked in 2% or 5% BSA milk or LI-COR TBS blocking buffer for 1 h. The membranes were then incubated with primary antibody overnight at 4° C. Membranes were washed with TBST (0.1-2% Tween 20) and TBS before and after incubation with secondary antibody (LI-COR, CST 7074S) for 1 h at room temperature. Membranes were imaged on the Odyssey CLx Imagining system (LI-COR) or iBright750 (Invitrogen) and analyzed using LI-COR ImageStudio or ImageJ. The following antibodies were purchased from Cell Signaling Technology: G9a (C6H3), GLP (E6Q8B), H3K9me2 (D85B4), vinculin (E1E9V), β-Actin (13E5).

### Cell viability assay

22Rv1, K562, H1299 and PNT2 cells were seeded in 96 well-plates and treated with compound at indicated serial dilations for 5-8 days. Plates were incubated at 37° C with 5% CO_2_. Cell viability was determined using CCK-8 (Cell Counting Kit-8, Dojindo, CK04). Absorbance and reference values were measured at 450 nm and 690 nm respectively on the Infinite F PLEX plate reader (TECAN). IC_50_ values were calculated using GraphPad Prism 6.

### Clonogenicity assay

22Rv1 cells were seeded at 50,000 cells per well in 6 well plates and treated continuously with compounds at indicated concentrations for 8 days. After 8 days, the cells were stained with 0.5% crystal violet dye.

### Methyltransferase inhibition assays

G9a and GLP enzymatic activity assays were conducted in duplicate at indicated concentrations using the radioisotope-based MT HotSpot™ system by Reaction Biology Corp. The selectivity panel of 21 methyltransferases was performed in duplicate with the same assay by Reaction Biology Corp. using 10 µM of compound.

### RT-qPCR

RT-qPCR was performed according to a previously described protocol ^3157^. H1299 cells were treated with DMSO, UNC0642 (3 µM), or MS8709 (3 µM) for 24 h. Total RNA was extracted using the Monarch Total RNA Miniprep Kit (T2010S, New England Biolabs) and cDNA was synthesized using the SuperScript IV First-Strand Synthesis System (18091050, Thermofisher). qPCR was performed on the Agilent Technologies Stratagene Mx3005p qPCR system. Primer sets used are listed below.

### Mouse PK studies

Compound **10** (in its HCl salt form) was dissolved in a solution formulation of 5% NMP, 5% Solutol HS-15, and 90% Normal saline. Three male C57BL/6 mice were administered intraperitoneally with solution formulation of compound **10** at 50 mg/kg. Blood samples (approximately 60 µL) were collected under light isoflurane anesthesia from a set of three mice at, 0.5, 2 and 8 h. Plasma was harvested by centrifugation of blood and stored at -70 ± 10 °C until analysis. Plasma samples were quantified by fit-for-purpose LC-MS/MS method (LLOQ: 5 ng/mL). Pharmacokinetic analysis was performed using GraphPad Prism software in a way of nonlinear regression analysis. Compound concentrations in plasma at each time point are the average values from 3 test mice. Error bars represent ± SEM. This PK study was conducted in compliance with Institutional Animal Care and Use Committee (IACUC) guidelines.

## ASSOCIATED CONTENT

### Supporting Information

The following Supporting Information (SI) is available free of charge on the ACS Publications website at DOI. Molecular formula strings for all compounds (CSV), Figures S1 to S3, Table S1, LC-MS, ^1^H NMR and ^13^C NMR spectra of compounds **10** and **14**.

### Author Contributions

_#_J.V. and Y.H. contributed equally to this work.

### Notes

J.J. is a cofounder and equity shareholder in Cullgen, Inc., a scientific cofounder and scientific advisory board member of Onsero Therapeutics, Inc., and a consultant for Cullgen, Inc., EpiCypher, Inc., Accent Therapeutics, Inc, and Tavotek Biotherapeutics, Inc. The Jin laboratory received research funds from Celgene Corporation, Levo Therapeutics, Inc., Cullgen, Inc. and Cullinan Oncology, Inc. Other authors declare no conflicts of interest.

## ACKNOWLEDGMENTS

This work was supported in part by the National Institutes of Health (NIH) grant P30CA196521 (to J.J.) and an endowed professorship from the Icahn School of Medicine at Mount Sinai (to J.J.). J.V. acknowledges the support by the Training Grant in Cancer Biology no. T32CA078207 at the Icahn School of Medicine at Mount Sinai from the National Cancer Institute of the NIH. This work utilized the NMR Spectrometer Systems at Mount Sinai acquired with funding from the NIH SIG grants 1S10OD025132 and 1S10OD028504.

## ABBREVIATIONS USED

DMF: dimethylformaldehyde
DIEA: di-isopropylethylamine
EHMT1: Euchromatic histone-lysine N-methyltransferase 1
EHMT2: Euchromatic histone-lysine N-methyltransferase 2
GLP: G9a like protein
HATU: hexafluorophosphate azabenzotriazole tetramethyl uranium
H3K9me2: histone H3 lysine 9 dimethylation
IP: intraperitoneal
MOA: mechanism of action
MW: microwave irradiation
NSCLC: non-small cell lung cancer
PK: pharmacolinetic
POI: protein of interest
PPIs: protein-protein interactions
PROTAC: proteolysis targeting chimera
SAR: structure-activity relationship
TPD: targeting protein degradation
UPS: ubiquitin proteasome system
VHL: von Hippel-lindau
WB: western blotting;

