## Supplemental figures for "Discovery of the First-in-class G9a/GLP PROTAC Degrader"

[2]Current address: College of Pharmacy, Keimyung University, Daegu 704-701, South Korea

### Table of Contents

**Figure S1.** Representative WB results of compounds **1-13** on reducing G9a and GLP protein levels **S3**

**Figure S2.** Compound **10** is non-toxic in the PNT2 normal human prostate cell line **S4**

**Figure S3.** The effect of compound **10** and UNC0642 on the mRNA levels of *G9a* and *GLP* in H1299 cells **S5**

**Table S1.** PK parameters of compound **10** in C57BL/6 mice following a single IP administration **S6**

$^{1}$H NMR spectrum of compound **10** **S7**

$^{13}$C NMR spectrum of compound **10** **S8**

LC-MS spectrum of compound **10** **S9**

$^{1}$H NMR spectrum of compound **14** **S10**

$^{13}$C NMR spectrum of compound **14** **S11**

LC-MS spectrum of compound **14** **S12**

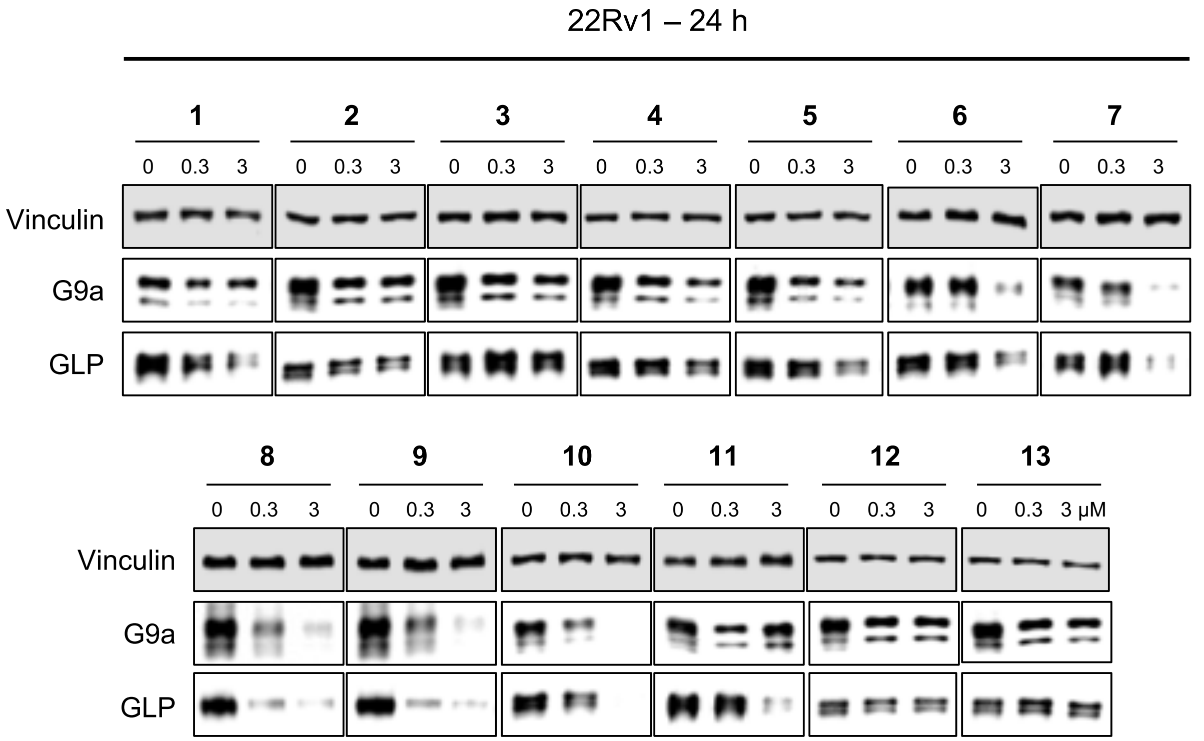

**Figure S1.** Representative WB results of compounds **1-13** on reducing G9a and GLP protein levels. 22Rv1 cells were treated with DMSO or the indicated compound at 0.3 or 3 μM for 24 h. The cell lysates were analyzed via WB to examine G9a and GLP protein levels with vinculin as the loading control. Results shown are representative from two independent experiments.

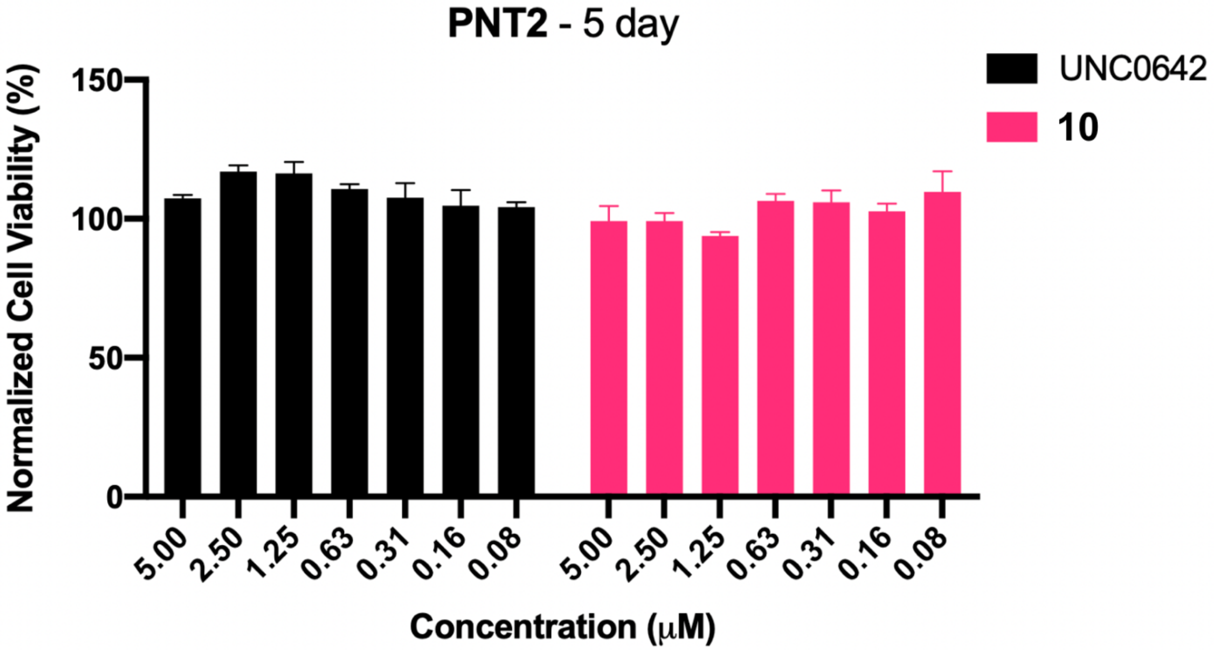


**Figure S2.** Compound **10** is non-toxic in the PNT2 normal human prostate cell line. PNT2 cells were treated with compound **10** or UNC0642 at the indicated concentration for 5 days in the WST-8 assay (CCK-8). Data shown are the mean values ± SD from 2 biological repeats.

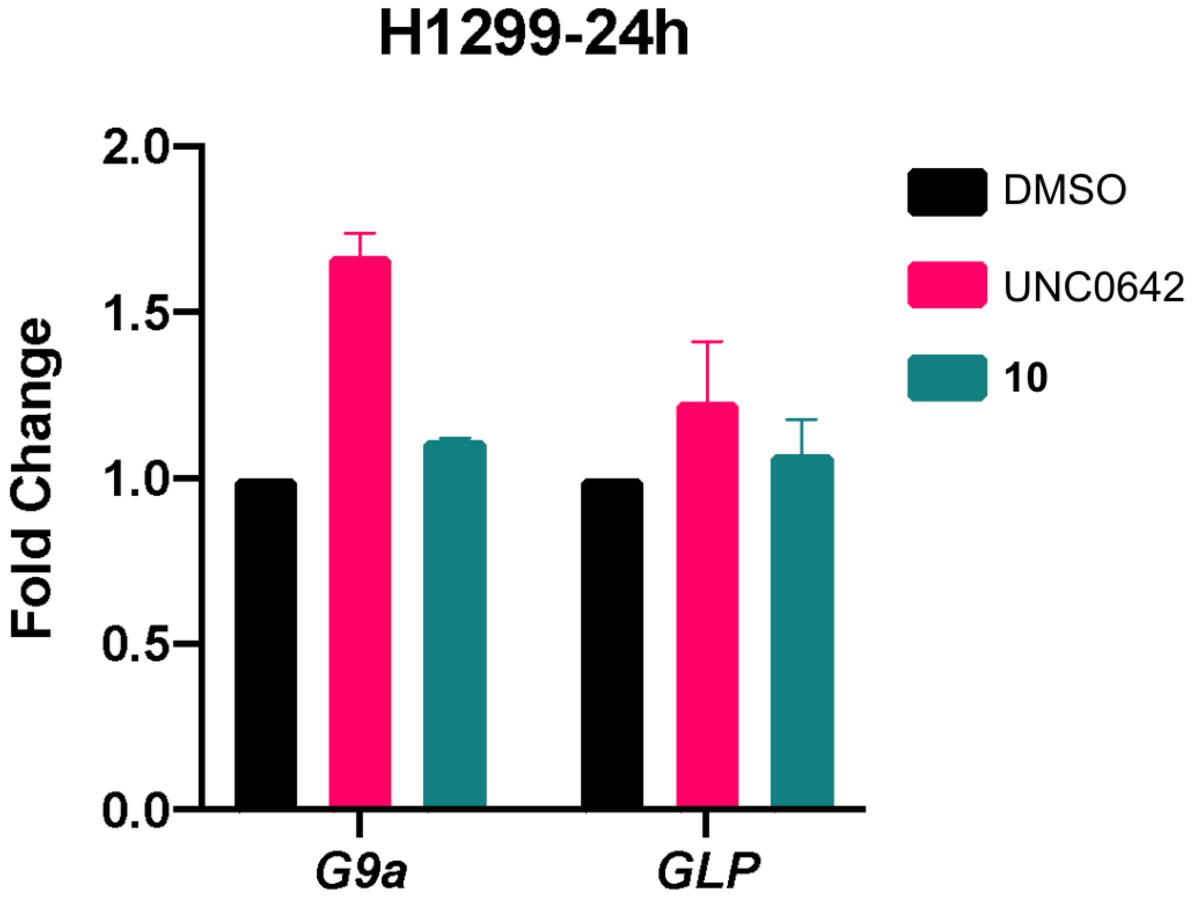


**Figure S3.** The effect of compound **10** and UNC0642 on the mRNA levels of *G9a* and *GLP* in H1299 cells. The mRNA levels of *G9a* and *GLP* were determined using RT-qPCR. H1299 cells were treated with DMSO or the indicated compound at 3 μM for 24 h. The mRNA levels were normalized to the DMSO control. The data shown represent the means ± SD from two independent experiments.

**Table S1.** PK parameters of compound **10** in C57BL/6 mice following a single IP administration.

| **Compound** | **Route** | **Dose (mg/kg)** | **Matrix** | **$T_{max}$ (h)** | **$C_{max}$ (μM)** | **$AUC_{last}$ (hr*μM)** | **$T_{1/2}$* (h)** |
|---|---|---|---|---|---|---|---|
| Compound **10** | IP | 50 | Plasma | 0.5 | 27 ± 0.6 | 61 ± 3 | 1.5 |

*Estimated result.

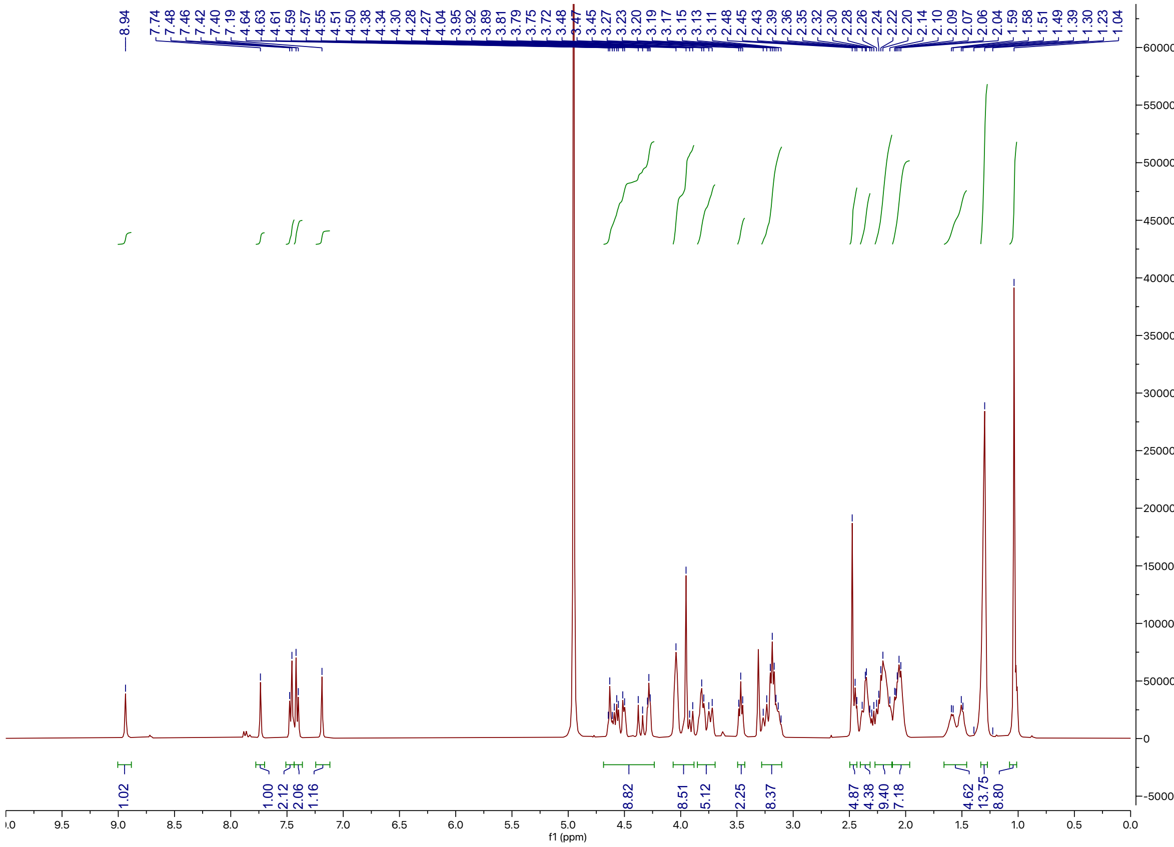

[1]H NMR spectrum of compound **10**

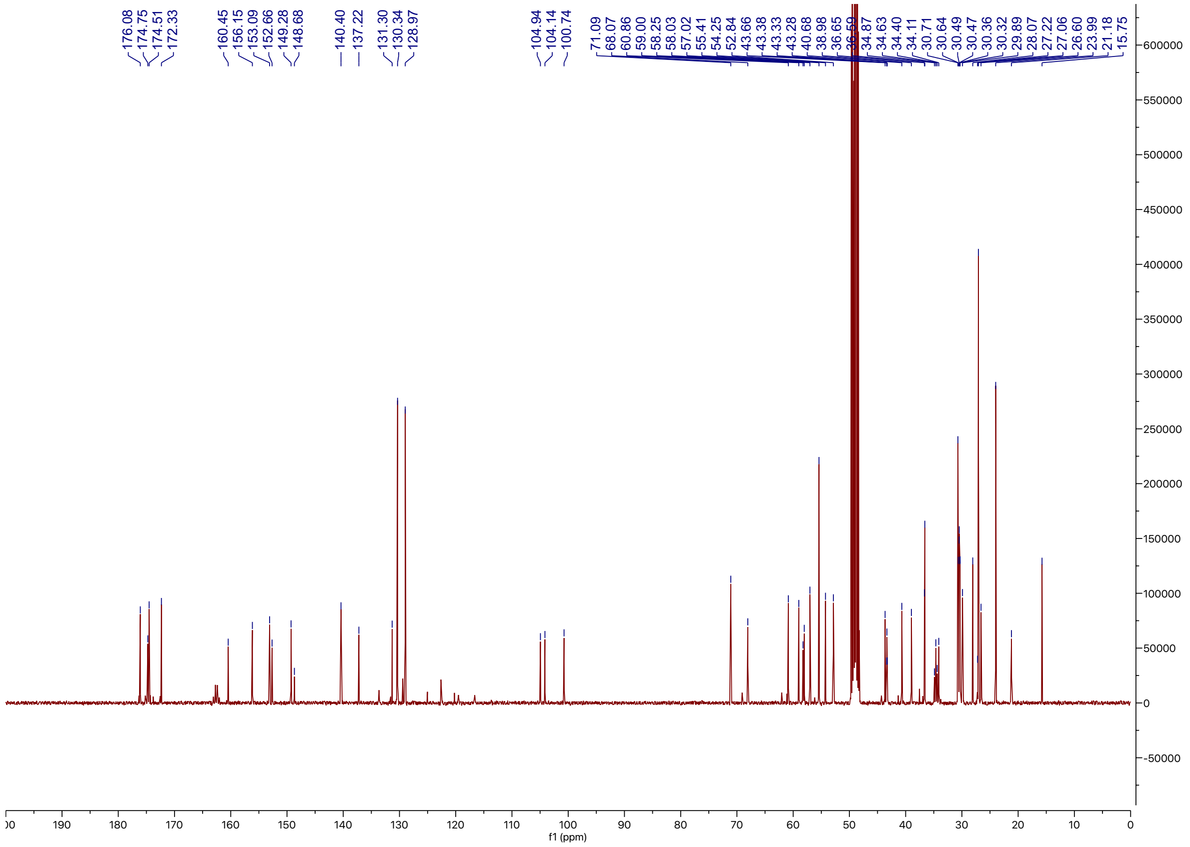

$^{13}$C NMR spectrum of compound **10**

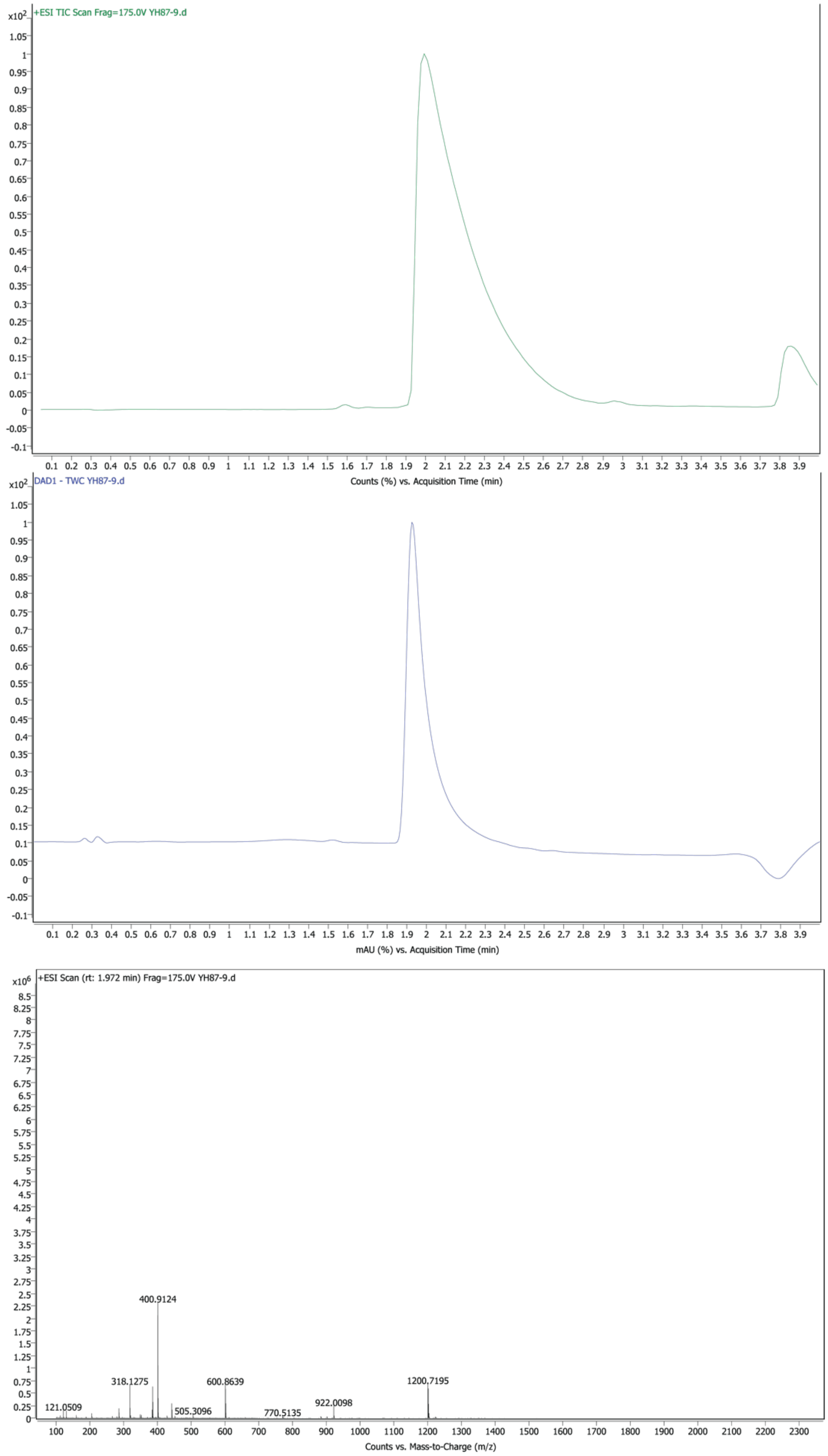


LC-MS spectrum of compound **10**

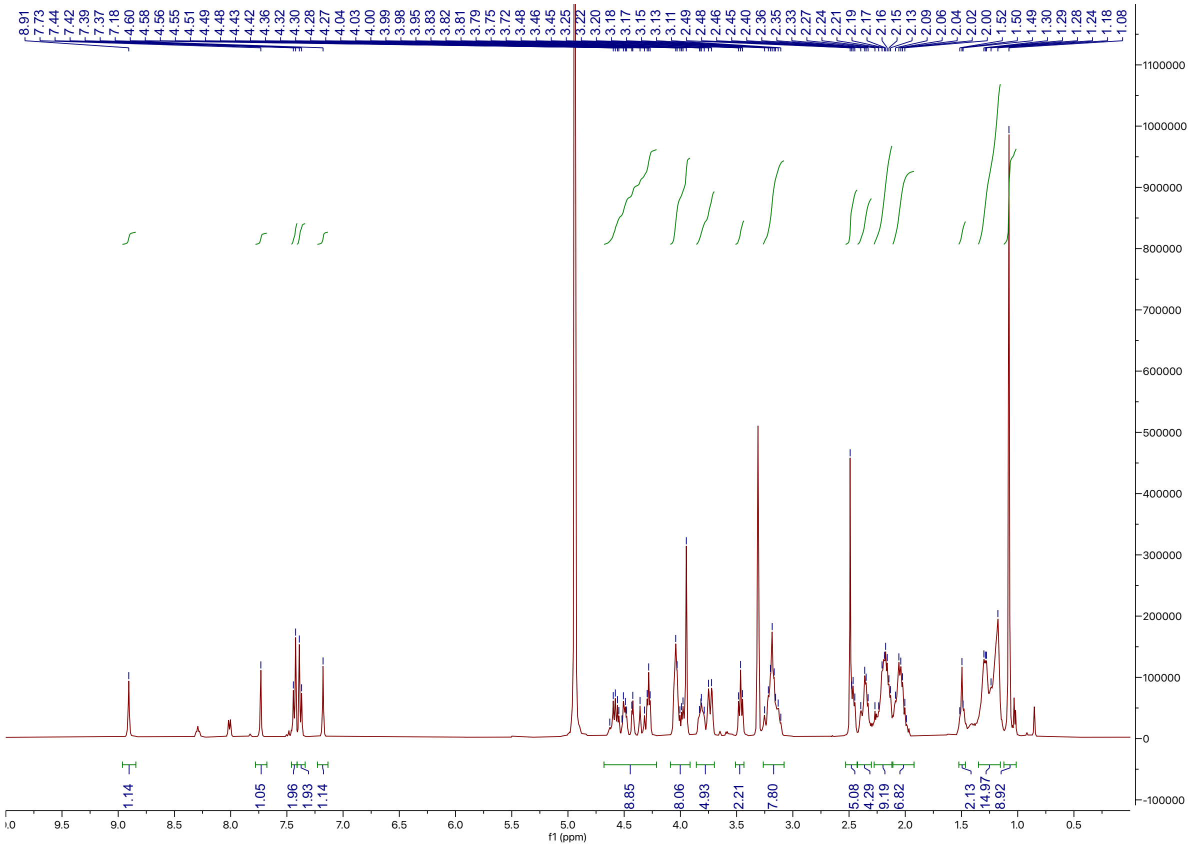

[1]H NMR spectrum of compound **14**

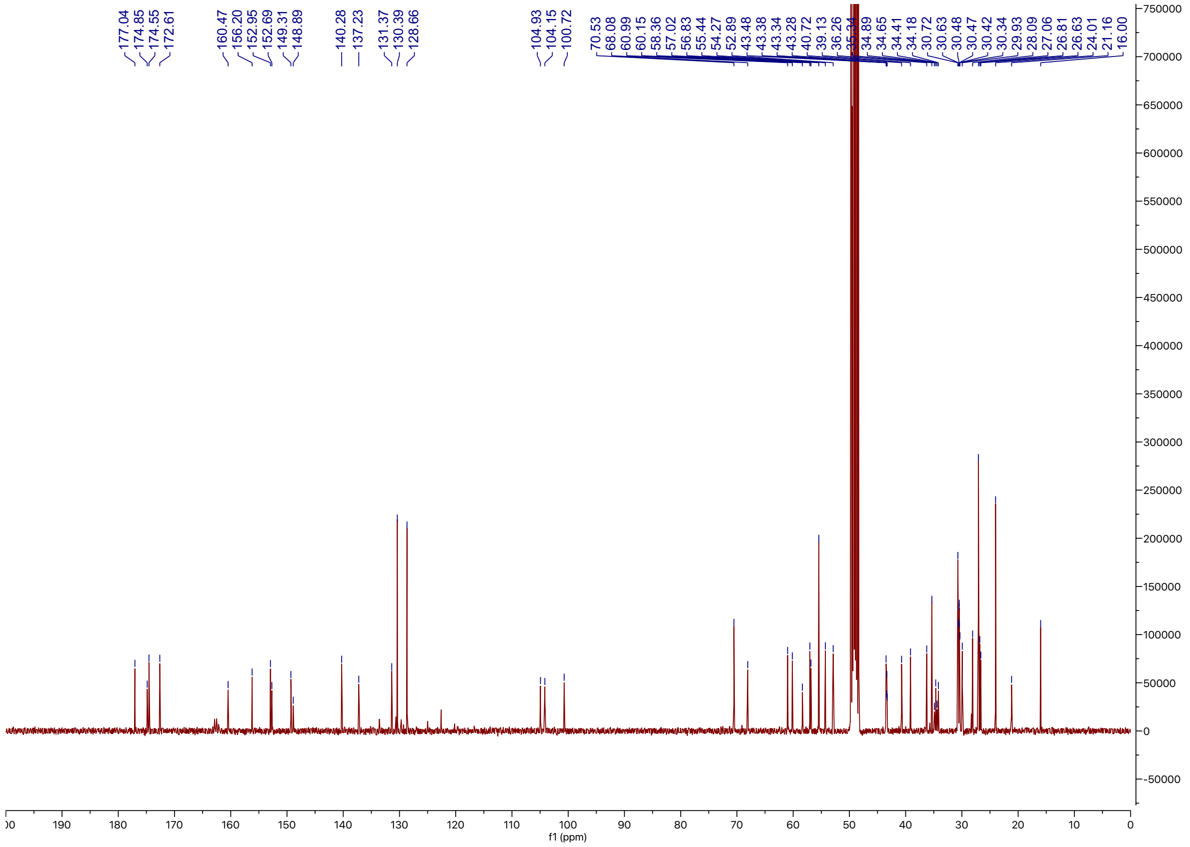


$^{13}$C NMR spectrum of compound **14**

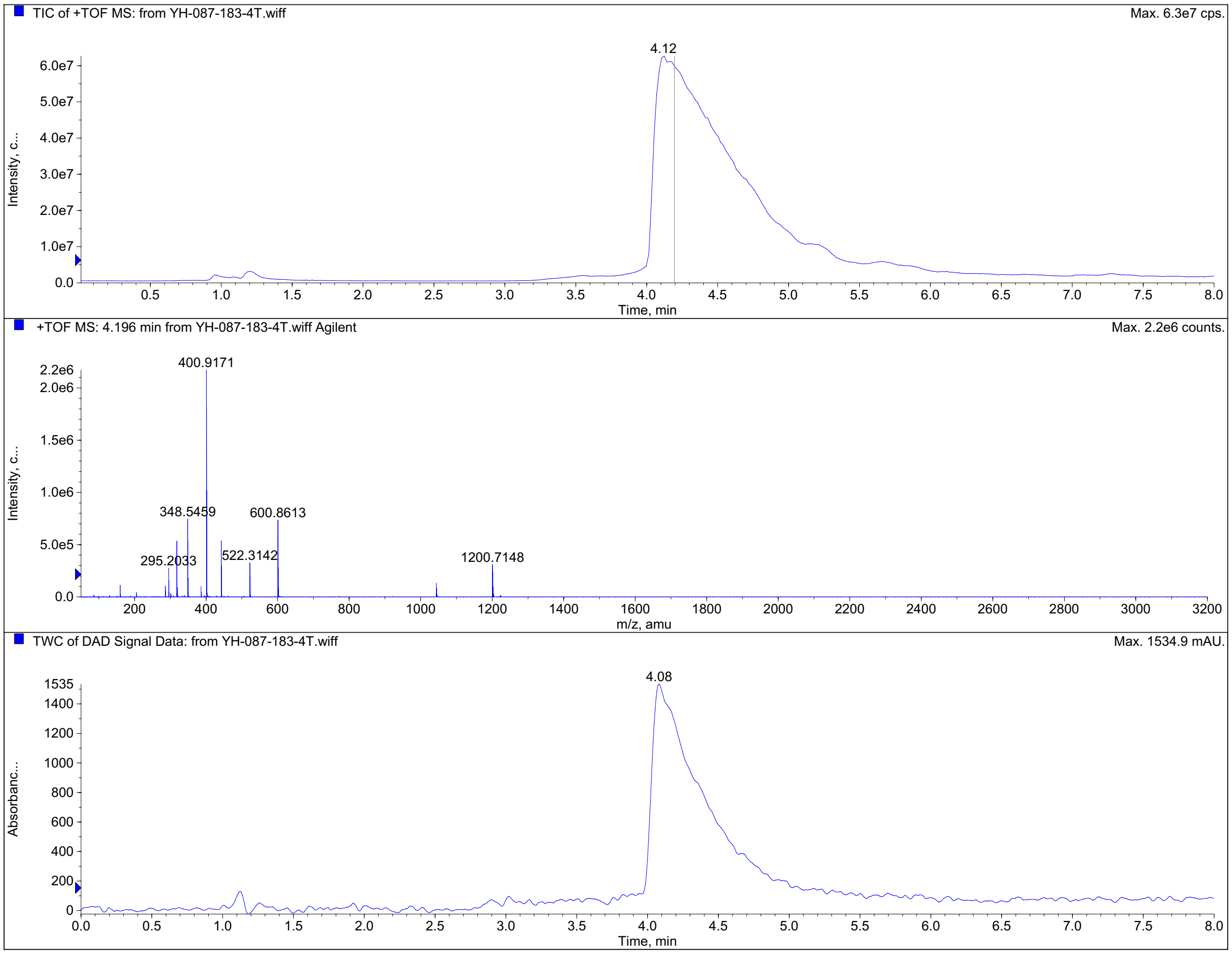


LC-MS spectrum of compound **14**
